# Regulatory mode and enhancer architecture determine limits of blood DNA methylation as a molecular proxy

**DOI:** 10.64898/2026.07.14.738581

**Authors:** Varun B. Dwaraka, Max Melnikas, Sayf Al-Deen Hassouneh, Ashley E. Rosko, Carolyn J. Presley, Andrea Aparicio, Ryan Smith, Jessica Lasky-Su, Christin E. Burd

## Abstract

Blood-based DNA methylation is widely used as a molecular surrogate for traits ranging from circulating protein and metabolite levels to biological aging, but whether the CpGs driving these models reflect genuine gene regulation or statistical convenience is unresolved. Training elastic-net models using the expression of 77 T cell genes across 333 matched T-cell enriched blood samples, we show that predictability is governed by regulatory mode: genes associated with differentiation and lineage-identity are strongly predictable, whereas dynamically regulated genes encoding cytokines and immune checkpoint ligands are not. The predictive CpGs are neither at the gene promoter nor at master-regulator binding sites; instead, the signal is carried by gene-specific sets of distal CpGs enriched within T-cell enhancer chromatin states distributed throughout the genome. Methylation-predicted gene expression validated against directly measured NanoString values in an independent 228-sample cancer cohort, remaining highly concordant for lineage genes but only weakly concordant for stimulus-responsive checkpoint ligands, confirming the lineage-versus-stimulus divide independently of the training cohort. On-gene retraining and chromatin-state annotation provide complementary tests of the genomic context of predictive CpGs.

## Introduction

Blood-based DNA methylation measures have enabled a growing class of predictive surrogates for molecular and clinical phenotypes that would otherwise require additional, and sometimes costly or invasive, measurements. DNAm GrimAge, for example, combines chronological age and sex with DNA methylation surrogates for smoking pack-years and seven plasma proteins to predict mortality risk and healthspan [1]. Its successor, GrimAge2, uses the same set of 1,030 CpGs while adding methylation surrogates for C-reactive protein and hemoglobin A1c [2]. This strategy has since been extended beyond individual aging clocks. Gadd et al. trained epigenetic scores, or EpiScores, for 953 circulating proteins, of which 109 met their validation criteria [3]. DNAmFitAge incorporates methylation surrogates for gait speed, grip strength, forced expiratory volume in one second, and maximal oxygen uptake [4], whereas PhysAge integrates methylation surrogates for eight biomarkers representing multiple physiological systems[5]. Composite frameworks such as OMICmAge further this idea, combining DNA methylation surrogates for proteomic, metabolomic, and clinical-laboratory values into a single aging biomarker [6]. Such proxy-based architectures are increasingly used to predict aging-related outcomes and disease risk and may offer greater interpretability than models defined only by weighted sets of CpGs.

Despite their predictive utility, the biological basis of these proxies remains incompletely understood. In most cases, CpGs are selected by penalized regression or other machine-learning methods to maximize out-of-sample prediction, without requiring that the selected loci have a regulatory relationship with the target phenotype. Two related questions therefore remain unresolved: whether the CpGs carrying the greatest predictive weight occur at genomic elements with regulatory relevance to the target, and whether such regulatory grounding explains why some molecular traits are substantially more predictable from DNA methylation than others. Expression quantitative trait methylation (eQTM) studies have identified extensive associations between DNA methylation patterns and gene expression, encompassing both promoter-proximal and distal effects and showing enrichment within regulatory regions [7, 8]. However, pairwise eQTM mapping does not establish whether a gene expression can be accurately inferred from the methylome, which CpG combinations contribute predictive information, or whether the regulatory context explains variation in predictive performance across genes.

We addressed these questions using paired gene expression and DNA methylation data because transcription represents a proximal downstream consequence of DNA methylation and provides a mechanistically interpretable framework for evaluating whether predictive CpGs reflect regulatory biology. Large-scale eQTM studies have established that DNA methylation and gene expression are extensively associated across the genome, but these relationships vary substantially across genes, genomic contexts, tissues, and cellular states [7, 8]. The conditions under which methylation reliably supports accurate expression prediction remain incompletely understood, and defining those conditions would provide a principled framework for interpreting methylation proxies for more distal molecular phenotypes, including circulating proteins and metabolites.

T cells provide a particularly informative model system for studying the relationship between DNA methylation and gene expression because their subsets and differentiation states are defined by well-characterized genomic and phenotypic programs. Lineage-associated genes such as *CD8A* and *RUNX3* are linked to stable, heritable DNA methylation patterns established during differentiation and maintained across circulating T cell subsets. Consistent with this, locus-specific DNA methylation has been shown to regulate both cytotoxic and regulatory T-cell identity [9, 10, 11]. In contrast, genes regulated by NF-kappaB, interferon-JAK-STAT, and related signaling pathways, including *IL6*, *CD274*/PD-L1, and *TNF*, typically exhibit more dynamic and context-dependent DNA methylation patterns that reflect the cell’s current signaling environment rather than its differentiation history [12, 13]. Examining this contrast across diverse T cell transcriptional programs may help distinguish methylation marks that are directly coupled to gene regulation versus those that reflect indirect or transient relationships.

First, if methylation-based prediction models derive from durable regulatory coupling rather than expression abundance or statistical convenience, lineage-associated genes should be more accurately predicted than stimulus-responsive genes. Second, the CpGs carrying predictive information should fall within T cell cis-regulatory elements, such as lineage-specific enhancers and promoters. Notably, how such regulatory elements are organized is unclear: predictive signal could be concentrated at the target gene itself, distributed across distal enhancers, or organized around shared lineage-regulator binding sites.

Using 333 samples with paired genome-wide DNA methylation and NanoString expression data for 77 T cell genes, we evaluate the extent to which gene expression can be inferred from DNA methylation patterns and whether differences in predictive accuracy are associated with lineage-associated versus stimulus-responsive biology rather than simply reflecting transcript abundance. We then identify the CpGs underlying the predictive performance and test whether they form local, shared, or distributed regulatory architectures. To determine whether these models capture transferable biology rather than cohort-specific associations, we evaluate them in an independent cohort and test their robustness when applied to mixed-cell PBMC methylation patterns. Finally, as a population-scale generalizability check, we apply admixture-aware models to an independent whole-blood biobank to confirm the predicted scores retain expected relationships with age.

## Materials and methods

### Ethics and study approval

All human-subjects procedures for the training and cancer-validation cohorts were approved by the Institutional Review Board of The Ohio State University (training cohort: OSU-2017H0338; cancer-validation cohort: OSU-2018C0069), and all participants provided written informed consent; the study was conducted in accordance with the Declaration of Helsinki. The external whole-blood cohort (Mass General Brigham Biobank) was analyzed under the MGB institutional data-access agreement. Data for the retrospective analyses were accessed for research purposes on 2 July 2026. The authors did not receive or have access to protected health information or other participant-identifying information.

### Cohorts and samples

Models were trained in the Ohio State Microbiome and Aging (MAG) cohort. From this resource, 333 samples with matched Illumina EPIC v1 DNA methylation, NanoString T cell panel expression, and phenotype data were selected; all participants had purified T cell samples at the time of methylation assay. Data were partitioned into training and held-out test sets using a subject-level 80/20 split (seed 42), so that all visits from the same subject were assigned to the same fold, precluding information leakage from longitudinal measurements.

The trained models were validated in an independent cancer cohort assembled by Burd and Rosko (Ohio State University Comprehensive Cancer Center), comprising 228 subjects: 29 healthy donors, 106 hematologic malignancies (including chronic lymphocytic leukemia, acute leukemia, lymphoma, plasma cell disorders, and other hematologic cancers), and 93 lung cancers. Of these, 191 had paired Illumina EPIC v1 methylation and NanoString expression data enabling direct concordance assessment; all 228 were used for prediction. Chronological age was available for 224 of 228 subjects. Per the cohort’s CD3-based reporter sample enrichment strategy, samples are enriched for T cells but are not FACS-sorted to purity; the degree of residual non-T-cell content, and whether it differs by diagnosis, was assessed empirically by reference-based deconvolution (see External cancer-cohort validation, below) rather than assumed. Healthy donors were substantially younger (mean age 51.8 years, SD 20.5, range 22–87) than every disease group (mean age 70.1 to 73.6 years across the six malignancy groups, SD approximately 6 to 7 years); this age difference is a design feature of comparing research-volunteer controls to a clinical cancer cohort and is addressed explicitly in the disease-association statistical models described below. A subset of patients contributed more than one methylation sample to this cohort (37 hematologic-malignancy patients with a paired baseline and follow-up draw, and 33 lung-cancer patients with 2–3 serial on-treatment draws over up to two years); disease-diagnosis association testing (below) is therefore restricted to one baseline, pre-treatment sample per patient (n = 131 of the 224 age-annotated subjects), whereas concordance assessment (Fig. 4a) and the per-sample data release (Supplementary Data 6) use the full available sample.

To test model robustness to cellular admixture, 385 peripheral blood mononuclear cell (PBMC) samples from the same MAG cohort participants were analyzed. These samples had matched purified T cell NanoString expression (362 paired) and were processed on Illumina EPIC v1 chips assayed in a single batch (206xxx chips).

For population-scale deployment, the PBMC model set was applied to the Mass General Brigham (MGB) Biobank whole-blood cohort (n = 4,250 with complete phenotype data for outcome analyses; 4,483 with age data for the validity gate; Illumina EPIC v1; ssNoob normalization). This cohort has electronic-health-record-linked longitudinal outcomes provided via the MGB Biobank infrastructure; disease incidence and all-cause mortality were ascertained from structured EHR data. The genome build was hg19 throughout.

### DNA methylation array processing and normalization

All cohorts were assayed on the Illumina Infinium MethylationEPIC v1 (EPIC) BeadChip. Raw IDAT files were processed in R 4.4.0 using the minfi package v1.52.1 [23].

The training (MAG T cell, n = 333) and PBMC (n = 385) datasets were processed together from IDAT files using reference-anchored functional normalization, with control-probe intensities projected into a pre-computed principal-component space (3 PCs, approximately 22,000 whole-blood EPIC v1 samples) to standardize across batches independent of the current sample’s size. Background/dye-bias correction and RCP-based probe-type-bias correction were then applied, and CpGs with detection p greater than 0.05 were masked to NA. Full parameters and rationale are in Supplementary Note 2.

The cancer cohort (Burd/Rosko) was normalized independently using functional normalization followed by regression on correlated probes (RCP[25]). RCP was applied to minimize residual technical variation between cancer and control samples processed across multiple scanner runs.

The MGB Biobank cohort was provided as a precomputed single-sample Noob (ssNoob) beta matrix [26], normalized independently of the training pipeline by the MGB team. Because ssNoob differs from the FunNorm plus RCP normalization used in training, the MGB analysis constitutes a cross-normalization replication; robustness was confirmed by the p16-versus-age positive control before any disease testing.

CpGs were annotated to gene, genomic region, and CpG-island context using the hg19 EPIC manifest (IlluminaHumanMethylationEPICanno.ilm10b4.hg19, Bioconductor v0.6.0). M-values (logit2-transformed beta values, bounded away from 0 and 1 at 0.001 and 0.999) were used for all model fitting and statistical testing because the heteroskedasticity of beta values violates linear model assumptions; beta values are used only for effect-size reporting and visualization.

### NanoString expression quantification

T cell gene expression was measured on the NanoString nCounter Human T Cell Characterization Panel (77 genes plus housekeeping controls), with upper-quartile normalization, RUVg-based unwanted-variation removal (k = 1), DESeq2 size-factor normalization, and log2 transformation. Two panel targets with more than 25% of samples at the background-expression floor (GRAIL/RNF128, VTCN1/B7-H4) were excluded, leaving 77 genes for modeling. Full normalization parameters are provided in Supplementary Note 2.

### Methylation-to-expression model training

For each of the 77 genes, candidate CpGs were identified genome-wide by methylation-expression association, selecting between Pearson, Spearman, and a limma duplicateCorrelation model (to accommodate repeated-measures structure) by cross-validation; the top 50,000 CpGs by association statistic were retained per gene. Full feature-selection methodology is provided in Supplementary Note 2.

Predictive models were elastic nets fit with glmnet v5.0, chosen for sparse feature selection under the collinearity inherent in genome-wide methylation data. Alpha was tuned over a six-value grid (near-ridge to lasso) with subject-level three-fold outer cross-validation (seed 42) and 10-fold inner cross-validation for lambda (lambda.min criterion); the final model was refit on all training data at the best alpha. Cross-validated R^2^ was floored at zero to prevent negative values from inflating apparent performance for uninformative genes. Full hyperparameter grid, fold-assignment, and refitting details are provided in Supplementary Note 2.

Stable CpGs were defined as those with a non-zero elastic-net coefficient in at least two of three outer folds. Genes were grouped into four interpretive performance tiers by cross-validated R^2^ (High at or above 0.70; Moderate 0.55 to 0.70; Weak 0.40 to 0.55; Uncoupled below 0.40); all quantitative analyses use R^2^ as a continuous variable. Tier-boundary rationale is provided in Supplementary Note 2.

### Localizing the predictive signal: on-gene versus off-gene audit

Every stable CpG was classified as on-gene (target gene’s own promoter/body, hg19 EPIC manifest) or off-gene (all other locations), and the weight share of each class was computed from summed absolute elastic-net coefficients. To test whether off-gene CpGs carry information beyond their number, an on-gene-restricted elastic net was retrained per gene and compared to the genome-wide model on the identical held-out split. Full retraining parameters and the on-gene-only comparison methodology are provided in Supplementary Note 2.

To test for shared upstream regulators, the top-10 highest-absolute-coefficient driver CpGs from each model were pooled across all 77 genes and their recurrence across two or more models was quantified; a strict variant further intersected recurring drivers with enhancer chromatin states (defined below). Host-gene GO enrichment for off-gene CpG pools was computed as described in the Functional Enrichment section.

Gene-by-gene shared-transcription-factor test. To test whether each gene’s own extragenic driver CpGs are organized around that gene’s own known regulators, rather than only asking whether driver CpGs recur across different genes’ models, we mapped each gene’s top 20 off-gene driver CpGs to neighboring genes within 500 kb and tested, for 45 genes with a curated TRRUST v2 regulator, whether neighboring genes shared a transcription factor with the index gene more often than expected (one-sided hypergeometric test). None of the 45 testable genes reached nominal significance (smallest p = 0.71). Full testability criteria and per-gene results are provided in Supplementary Note 2 and Supplementary Data S4.

Because TRRUST v2 curation is sparse for T cell lineage genes, we repeated this test for CD8A and RUNX3 using ChIP-Atlas ChIP-seq peak overlap as an independent evidence source, then extended it across all 77 genes and four available transcription factors (308 tests); 8 of 308 reached nominal significance, four of which involve a transcription factor with an unusually broad genome-wide binding background. Neither this test nor the TRRUST-based test found evidence that off-gene driver CpGs are organized around a small set of shared or canonical regulators. Full statistical details and caveats are provided in Supplementary Note 2 and Supplementary Data S4.

### Chromatin-state annotation

Stable CpGs from each model were annotated with the Roadmap Epigenomics 15-state ChromHMM core-marks mnemonic segmentations (hg19)[33] for resting primary T cells (E043 CD4 naive, E044 CD4 memory primary, E047 CD8 naive) and for PMA-ionomycin-activated primary T helper cells (E041 stimulated Th, E042 stimulated Th17), obtained from the Roadmap portal (egg2.wustl.edu). Genomic overlaps were computed with GenomicRanges v1.58.0[34] and rtracklayer v1.66.0. The enhancer state comprised ChromHMM states labeled Enh (strong enhancer) and EnhG (genic enhancer) in the 15-state model. The fraction of each gene’s stable CpGs falling in enhancer states was related to cross-validated R^2^ by Pearson correlation to test whether T cell enhancer localization predicts model performance.

### Transcription-factor binding-site overlap

Overlap with broadly active canonical transcription factors used ENCODE narrowPeak ChIP-seq for CTCF (CD4 naive T cells, ENCFF858TLX; CD8 naive T cells, ENCFF758CQW), ETS1 (GM12878, ENCFF332PGQ), and SP1 (K562, ENCFF076YZO); peaks were provided in GRCh38 and were lifted to hg19 using UCSC liftOver (hg38ToHg19.over.chain).

For lineage master regulators, pre-called hg19 ChIP-seq peaks were obtained from ChIP-Atlas (chip-atlas.dbcls.jp; stringency q less than 1 × 10^-5^)[35] for TBX21 (primary in vitro-differentiated Th1 cells, the only primary T cell dataset available), RUNX3 (NK-92 cell line), PRDM1 (U-266 myeloma line), and FOXO1 (germinal-center B cells); EOMES had no ChIP-seq available in any immune cell type across ChIP-Atlas, ENCODE, and ReMap [36] and was therefore excluded from the binding-site analysis.

Because predictive CpGs are enriched in open-sea and distal contexts whereas most TF peaks occur at CpG-island promoters, a naive overlap fraction confounds CpG-island compartment with binding-site depletion. To disentangle these effects, overlap was tested using a Cochran-Mantel-Haenszel (CMH) common odds ratio stratified by CpG-island context (Island, Shore, Shelf, OpenSea), with Robins-Breslow-Greenland 95% confidence intervals. The CMH stratification tests whether off-gene driver CpGs are depleted at TF peaks within each compartment separately, eliminating the compositional artifact. All genomic interval operations used GenomicRanges and rtracklayer.

### Functional enrichment

Tier-stratified functional enrichment compared the top and bottom cross-validated-R^2^ terciles (balanced to equal sample sizes for power; tercile boundaries were data-adaptive) as the foreground versus background set. The enrichment approach was gene-set overrepresentation (Fisher exact test with BH-FDR correction) rather than GSEA, because the 77-gene panel is too small for reliable rank-based enrichment statistics.

Enrichment was computed across seven annotation databases (Gene Ontology BP/MF/CC, KEGG, Reactome, and the MSigDB Hallmark and C7 immunologic collections) using clusterProfiler v4.14.6, with missMethyl::gometh v1.40.3 additionally applied where CpG-number bias against an EPIC-array background was a concern. P-values were Benjamini-Hochberg corrected within each collection independently; results across collections were not further combined. Full package versions, database parameters, and a citation note on the gometh methodology are provided in Supplementary Note 2.

### Regulatory-network analysis

Each gene’s transcriptional regulatory context was characterized using TRRUST v2. Regulatory out-degree (downstream curated targets, a proxy for master-regulator status) and in-degree (upstream TF regulators) were related to cross-validated R^2^ by Pearson correlation; genes absent from TRRUST v2 were assigned a degree of zero. An exploratory STRING protein-protein-interaction-degree analysis found no association and is reported with its caveats in Supplementary Note 2.

### External cancer-cohort validation

The 77 trained T cell models (one per gene, selected by cross-validated R^2^) were applied without retraining to the Burd/Rosko cancer cohort (N = 228). Methylation-predicted expression was compared to directly measured NanoString expression per gene via Pearson correlation (cross-cohort concordance, n = 191 paired samples). Group-level concordance was assessed as the Pearson correlation between diagnosis-group means of predicted and measured expression (up to r = 0.97 for CD8A); the variance ratio (predicted variance divided by measured variance, expected to be less than 1 when methylation captures stable between-subject but not within-diagnosis stochastic variation) was computed per gene.

Blood cell composition and the CD3-enrichment calibration. Blood cell composition in the cancer cohort was estimated by EpiDISH v2.22.0 using robust partial correlation with the cent12CT reference matrix (12 blood cell types; full panel listed in Supplementary Note 2).

To confirm the cancer cohort matched the model’s training context, sorted-T training samples deconvolved to approximately 83% T cell fraction under this reference, which served as a calibration baseline. Four of six cancer-cohort diagnosis groups matched this baseline closely; chronic lymphocytic leukemia and acute leukemia deviated (lower T cell, higher B cell fraction), consistent with residual circulating malignant cells surviving CD3 enrichment. This diagnosis-specific composition, not a general whole-blood composite, is what the adjustment below controls for. Full per-group values are provided in Supplementary Note 2.

#### Disease-diagnosis association testing, with the age confound identified and corrected

For p16/CDKN2A, separation across diagnoses was tested by Kruskal-Wallis; each diagnosis was compared individually to healthy donors by two-sided Wilcoxon rank-sum test. In this unadjusted comparison, predicted p16 was nominally higher in three of five evaluable disease groups relative to healthy donors. However, because healthy donors in this cohort are substantially younger than every disease group (Cohorts and samples, above) and predicted p16 independently correlates with chronological age across the cohort (Pearson r = 0.385, p = 5.7 × 10^-6^, n = 131 baseline samples), we tested whether these unadjusted diagnosis comparisons could be explained by age rather than disease status.

For p16/CDKN2A and seven additional genes selected a priori as biologically relevant comparators (CD8A, CD3E, GZMB, FOXP3, RUNX3, CD274, PDCD1), each diagnosis-versus-healthy effect was tested under three nested linear models of increasing adjustment, fit separately for each of the 5 diagnosis groups (n = 131 with complete age data, restricted to one baseline sample per patient):

1. Unadjusted: predicted_expression ∼ diagnosis
2. Age-adjusted: predicted_expression ∼ diagnosis + chronological_age
3. Fully adjusted: predicted_expression ∼ diagnosis + chronological_age + CD8_total + CD4_total + Treg + B_total + NK + Mono + Neu

where CD8_total and CD4_total are the sums of the naive and memory subsets of each lineage from the 12-cell cent12CT deconvolution, and Treg, B_total, NK, Mono, and Neu are as returned by EpiDISH; eosinophils and basophils were excluded for the rank-deficiency reason given in Supplementary Note 2, yielding 7 grouped covariates rather than the full 12 raw cent12CT columns.

An association was classified as surviving age adjustment if the diagnosis coefficient remained nominally significant (p < 0.05) in model 2 with the same sign as in model 1, and as surviving full adjustment if it additionally remained so in model 3, fit on n = 131 baseline samples (a diagnosis category present only in the pooled, multi-visit sample, “other hematologic malignancies,” was excluded as it does not correspond to any baseline patient). Of 12 gene-diagnosis pairs nominally significant in the unadjusted model (40 tests: 8 genes times 5 diagnosis groups), all three p16/CDKN2A associations lost significance after age adjustment; 4 of the remaining 9 pairs survived both age and full adjustment (CD8A, CD3E, RUNX3, and CD274, all in acute leukemia, all reflecting reduced predicted expression relative to healthy donors). These 40 tests were not corrected for multiple testing, as the smallest diagnosis groups (n = 5–14) make this an underpowered, hypothesis-generating screen rather than a confirmatory test; the full three-model comparison for every test is reported directly (Supplementary Data S7) rather than relying on a corrected p-value alone. Full rationale is provided in Supplementary Note 2. Complete per-gene, per-diagnosis results under all three models are provided in Supplementary Data S7.

Detectability and cell-fraction mechanism test (coupling shift). As a complementary test of whether weakly predictable genes (cross-validated R^2^ < 0.45) are truly uncoupled from methylation rather than limited by low expression variance and cell-fraction dilution, we compared each gene’s CpG-expression coupling strength in healthy donors to its highest-expression cancer-cohort diagnosis group among 15 candidate genes. Coupling strengthened in the higher-expression group for 12 of 15 genes (exact binomial p = 0.035). Full selection criteria and per-gene results are provided in Supplementary Note 2 and Supplementary Data S8.

### PBMC admixture transfer

T cell models were applied to whole-PBMC methylation from 385 MAG cohort PBMC samples, yielding a per-gene PBMC transfer correlation against matched purified T cell NanoString expression (n = 362 paired samples, all 77 genes). To distinguish a genuine methylation identity signal from a cell-fraction readout, partial correlations were computed controlling for PBMC T cell fraction and then for the same 7 grouped cell-fraction covariates used in the cancer-cohort analysis; signal surviving full adjustment is intrinsic to methylation rather than composition. Full covariate definitions are provided in Supplementary Note 2.

As an independent, exploratory test, admixture-aware models were retrained directly on PBMC methylation for a representative 12-gene panel spanning high-predictability lineage markers, composition-sensitive genes, and mid/low-predictability comparators, using a simplified single-alpha, single-split procedure rather than the primary models’ full grid. PBMC-retrained and T cell models were compared on matched held-out performance metrics; CpG overlap between the two was low (median approximately 1.5%). Full gene panel, retraining parameters, and rationale are provided in Supplementary Note 2.

### Whole-blood population-scale deployment

To demonstrate deployment at population scale, both T cell- and PBMC-weighted model sets were scored on the MGB/Harvard Biobank whole-blood cohort (Illumina EPIC v1, ssNoob; n = 4,250), with each predicted score standardized to mean 0/unit SD so hazard/odds ratios are per-SD effects. CD80 was excluded from the PBMC model only (near-zero variance) and retained in the T cell model (76 and 77 evaluable scores). Composition was estimated by EpiDISH RPC (cent12CT reference); disease models include 11 of 12 raw cell fractions as covariates. This covariate scheme and its difference from the 7-grouped scheme above are detailed in Supplementary Note 2.

Mandatory composition adjustment was applied to every result reported from this cohort, because whole blood is granulocyte-dominated (approximately 60% neutrophils) and T cells comprise only approximately 15 to 25% of white cells; without adjustment, any association with immune or age-related disease can be wholly explained by cell-fraction differences across individuals.

As a validity gate applied before any disease analysis, predicted p16/CDKN2A was regressed on chronological age both marginally and after composition adjustment (linear regression, age as continuous predictor), and the naive and CD8 lineage scores (CD27, CD28, CD8A, CD127, CD3E) were checked for the expected age-related decline. This gate was required to pass before any disease analysis was interpreted.

Disease outcome data were obtained from structured EHR records linked to the MGB Biobank. Incident outcomes were defined as a new ICD-coded diagnosis after methylation collection, excluding participants with a prevalent diagnosis at baseline; all-cause mortality was ascertained from hospital records and death registration. Twelve incident disease outcomes were tested, selected by the MGB collaborators (M.M., J.L.S.) based on availability and event count; no primary outcome was pre-specified, and the MGB analysis is treated as discovery-and-illustration rather than confirmatory. The full outcome list is provided in Supplementary Note 2.

For all-cause mortality and each incident disease, composition-adjusted Cox proportional-hazards models were fit using the survival package v3.7.0, chosen for standard handling of right-censored biobank follow-up and per-SD-interpretable hazard ratios; the proportional-hazards assumption was not formally tested at this scale (912 PBMC and 924 T cell models). Models were fit unadjusted, semi-adjusted (age, sex), and fully adjusted (age, sex, 11 cell fractions); only fully adjusted results are reported in the main text. Cross-sectional associations used logistic regression with the same covariate structure. Full model specifications are provided in Supplementary Note 2.

Benjamini-Hochberg FDR correction (threshold FDR less than 0.05) was applied separately within each model set: 912 tests (76 genes times 12 diseases) for the PBMC-weighted model, 924 (77 times 12) for the T cell-weighted model, plus separate mortality corrections (76 and 77 tests, respectively). No correction was applied across the two model sets, which were treated as parallel discovery analyses. Full test-family definitions are provided in Supplementary Note 2.

To characterize concordance between the two model sets, Pearson correlation of log(OR) was computed across all 912 gene-disease pairs for the 76 genes common to both. Low overall concordance (r = 0.01) indicates complementary rather than redundant readouts, consistent with the 1.5% CpG overlap found in the admixture analysis. Full comparison methodology is provided in Supplementary Note 2.

### Statistics, software, and reproducibility

All statistical tests are two-sided unless otherwise stated. Correlations are Pearson unless specified as Spearman. Effect sizes are reported with 95% confidence intervals where applicable. Multiple testing was controlled by Benjamini-Hochberg FDR correction applied within each defined analysis family: within each functional-enrichment database collection, within the MGB incidence scan (912 PBMC tests and 924 T cell tests), and within the MGB mortality analysis (76 PBMC tests and 77 T cell tests). BH-FDR was chosen over Bonferroni throughout because the tests within each family are positively correlated (genes share CpGs; diseases share participants), and BH-FDR controls the expected proportion of false discoveries rather than the probability of any false discovery, which is the appropriate operating characteristic for discovery analyses.

The following analysis families were deliberately left uncorrected for multiple testing, and this is stated explicitly for each rather than left implicit:

- The cancer-cohort p16 Kruskal-Wallis and per-diagnosis Wilcoxon tests (up to 6 comparisons), and the 48-test (8 genes by 6 diagnoses, under 3 nested models) disease-association family in the External cancer-cohort validation section: both are exploratory, hypothesis-generating screens in a cohort with diagnosis groups as small as n = 5, where FDR correction would not materially change interpretation and where the three-model (raw/age-adjusted/fully-adjusted) comparison itself is the primary robustness check reported.
- The coupling-shift binomial test (12 of 15 genes, p = 0.035): a single pre-specified test, not a family.

No correction was applied across distinct analysis families (for example, between the enrichment collections and the cancer-cohort disease-association tests). Analyses designated as exploratory (the cross-sectional cancer-diagnosis comparisons, the coupling-shift mechanism test, and the whole-blood disease deployment) are labeled as such throughout; confirmatory inference was not claimed for any of these analyses.

Analyses were performed in R 4.4.0, primarily in a spring-2026 environment with a later snapshot (2026-07-04) used only for the age-confound re-analysis and Figure 4 reconstruction, which did not require re-fitting any statistical model. Complete package versions, citations, and installation commands for both snapshots are provided in Supplementary Note 1.

Random seeds were fixed throughout (training/test split seed 42; final model refit seed 1234; PBMC-retrain single split seed 42). All external dataset accessions, genome builds, download dates, and cell-type deconvolution reference details are also provided in Supplementary Note 1 (deposited with the Zenodo code archive as DATA_SOURCES.md).

## Results

### Regulatory stability, not transcript abundance, predicts methylation-to-expression coupling

If DNA methylation stably records T cell lineage identity, it should predict genes whose expression is fixed during differentiation far better than genes whose expression is dictated by transient signaling. We tested this directly using a randomly selected training set representing 80% of the cohort (n=270). Using elastic net models, we selected CpGs whose methylation co-varies with each of the 77 transcripts measured by Nanostring and combined their weighted methylation values into a predicted expression score. Model accuracy was then evaluated on a held-out test set comprising the remaining 20% of samples initially excluded (n=63). Across 333 matched blood samples, cross-validated R_2_ ranged from 0.14 (*PDCD1LG2*) to 0.82 (*CD8A*) with a median of 0.47. 34 of 77 genes reached an R^2^ of at least 0.50 and 6 reached at least 0.70, many of which were markers of T cell differentiation or lineage commitment (*CD8A* = 0.82, *SPI1* = 0.80, *KLRG1* = 0.71, *CD3E* = 0.71, *CD27* = 0.71, and *RUNX3* = 0.70, **Fig. S1**). Among the weakest performers were inducible immune checkpoint ligands and cytokines (*PDCD1LG2/PD-L2* = 0.14, *CD274/PD-L1* = 0.21, IL-17A = 0.30, *IL-21* = 0.36 and *TGFB1 = 0.38*). Because the MAG cohort includes participants with longitudinal samples, we evaluated three feature-selection strategies that differed in how they handle repeated measurements; for 52 of 77 genes, the approach explicitly modeling within-subject correlation yielded the strongest predictive models.

Differences in model performance were largely independent of transcript abundance as mean NanoString expression showed only a weak correlation with cross-validated R^2^ (Spearman rho = +0.23, p = 0.049, **Fig. S2**). A subset of genes amplified prior to Nanostring analysis to improve detection (*p16/CDKN2A*, *ARF*, *IL6*, *IL17A*, and *FOXO1*) had lower R^2^ values (median 0.34 vs. 0.48, Wilcoxon p = 0.0017), but were not systematically lower in abundance (p = 0.30). Several genes, including *IL6* and *ARF*, were readily detectable across samples yet remained poorly predicted, suggesting that their weak coupling reflects stimulus-responsive regulation rather than a detection-floor artifact (**Methods**; **Supplementary Data 1**).

### Methylation-based prediction draws overwhelmingly from extragenic CpGs

To determine where each gene’s predictive signal resides, we first examined whether the strength of methylation-expression coupling explains differences in model performance across genes. The strength of stable CpG methylation-expression correlations across a gene was the dominant predictor of model performance (r = +0.80, R^2^ = 0.64, **Fig. 1a**). Genes whose stable CpGs showed stronger methylation-expression correlations were consistently predicted with greater accuracy, explaining 64% of the variance in cross-validated R^2^ across the gene panel.

**Figure 1.**
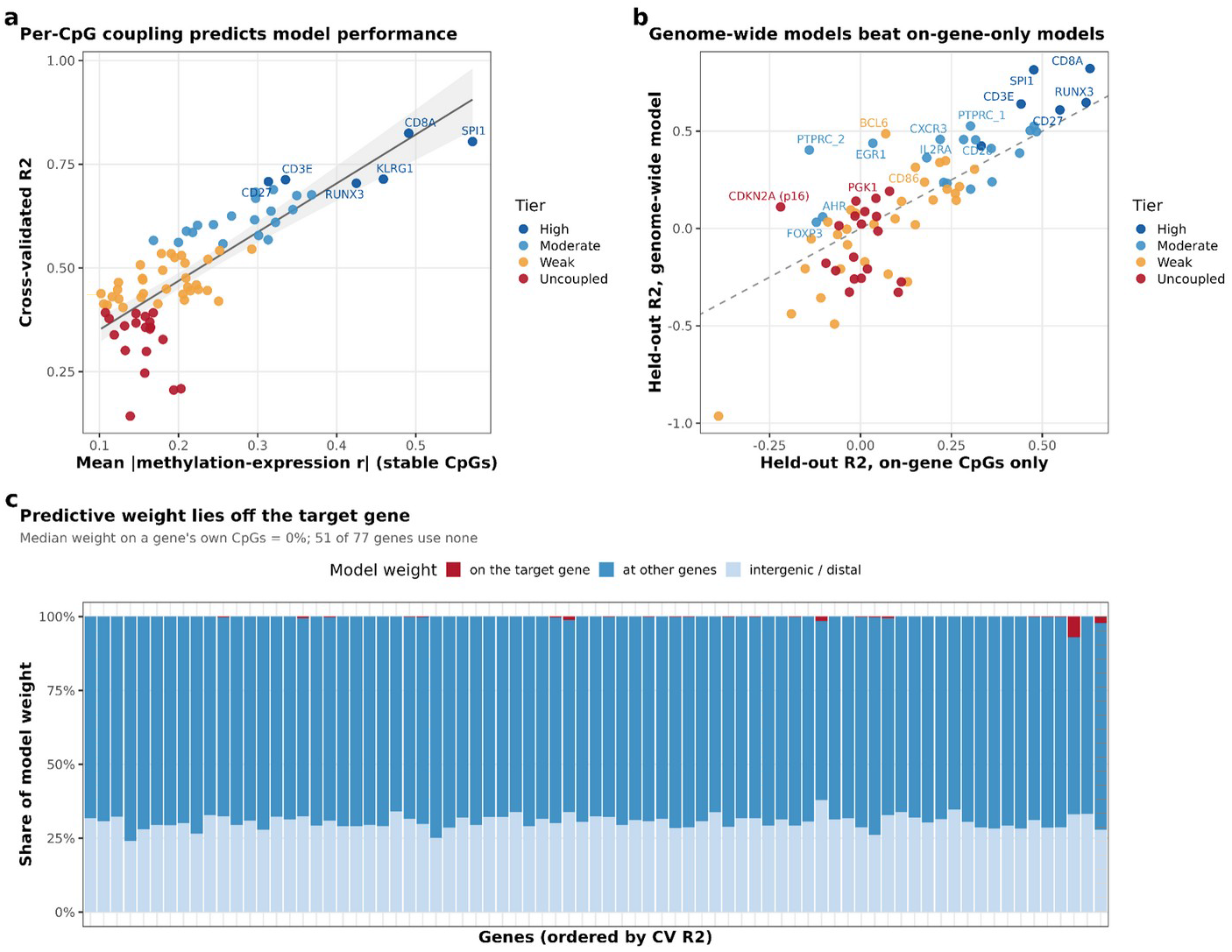
Blood DNA methylation predicts T cell lineage gene expression through distributed, off-gene CpGs. **A)**, Per-CpG methylation-expression coupling strength versus cross-validated R^2^ across 77 T cell genes. Each point is one gene; color indicates performance tier. The mean absolute methylation-expression correlation of a gene’s stable CpGs (x-axis) explained 64% of the variance in cross-validated R^2^ (r = 0.80, n = 77 genes). Labeled genes are discussed in the text. **B)**, Head-to-head comparison of genome-wide elastic net models versus models restricted to each gene’s own annotated CpGs (on-gene-only models), evaluated on the same held-out 20% test split. Each point is one gene; the dashed line is identity. Points below the diagonal indicate that restricting to on-gene CpGs degrades accuracy. Selected genes are labeled. **C)**, Share of total absolute model weight carried by CpGs annotated to the target gene itself (red), other genes (blue), or intergenic and distal sites (light blue), for each of the 77 genes ordered by cross-validated R^2^ (highest left). A total of 51 of 77 genes carry no on-gene weight. 333 matched blood samples (training); 20% held-out test set for panel b.

To directly test whether extragenic CpGs contribute predictive information beyond that contained within target gene CpGs, rather than simply reflecting a larger feature pool, we retrained elastic nets using only CpGs annotated to each target gene and compared their performance against genome-wide models on the identical held-out test set, though this on-gene comparison used a fixed hyperparameter rather than a matched optimization (Supplementary Note 2.18). Restricting to gene-annotated CpGs decreased accuracy for the majority of well-predicted genes: R^2^ declined from 0.82 to 0.48 for SPI1, from 0.64 to 0.44 for CD3E, and from 0.82 to 0.63 for CD8A. Some genes were nearly unpredictable from their own CpGs alone, including BCL6 (0.49 to 0.07) and EGR1 (0.44 to 0.03). A minority of lineage-associated genes showed little change, including RUNX3 (0.62 to 0.65), TBX21 (0.48 to 0.50), and CD3D (0.47 to 0.50), indicating that for this small subset, on-gene CpGs are sufficient predictors (**Fig. 1b; Supplementary Data 2**).

Consistent with this restriction result, genome-wide models had access to on-gene CpGs yet rarely selected them, arguing against a simple probe-count artifact: we examined the distribution of elastic net model weights across CpGs annotated to the target gene versus those located elsewhere in the genome. The median share of model weight attributable to CpGs annotated to the target gene was negligible, with 51 of 77 genes assigning no weight to their own CpGs whatsoever. Furthermore, no model placed more than 20% of its weight on gene-annotated CpGs, with the largest contributions observed for KLRG1 (7%) and CD8A (2.2%) (**Fig. 1c**). This distribution indicated that predictive signal derives predominantly from extragenic (outside of the annotated gene locus) rather than intragenic CpGs.

### The off-gene signal is gene-specific, not a shared regulatory hub

We next asked whether the extragenic CpGs contributing to prediction converged on shared regulatory mechanisms. If co-regulated gene programs shared predictive information through common regulatory elements, such as a T-bet or EOMES binding site contributing to multiple cytotoxic T cell gene models, we would expect their top driver CpGs to recur across models. We pooled the top 10 weighted distal CpG drivers for each of the 77 genes and tested performance. Pooling yielded 767 driver-CpG instances corresponding to 717 distinct CpGs; 672 (94%) appeared in only one gene’s model, and the remaining 45 (6%) recurred across two or more models, of which only 3 were in enhancer sites and none annotated to a canonical lineage master regulator. Across all 77 models, only 68 of 196,857 stable CpGs (0.03%) were annotated to the target gene itself; 51 of 77 used zero on-gene CpGs. The predictive signal is carried almost entirely by CpGs at other genes’ regulatory regions or intergenic and distal sites. The five genes with the largest on-gene contribution were *KLRG1* (2 of 186 total CpGs on-gene, carrying 7.1% of the total model weight), *CD8A* (11 of 1,682, 2.2%), *CCR6* (1 of 122, 1.4%), KIR Inhibiting Subgroup 1 (2 of 273, 1.1%), and *TIGIT* (5 of 2,564, 0.5%); even at these maxima, on-gene CpGs remain a small minority both by count (0.2-1.1% of each gene’s total CpGs) and by weight) (**Supplementary Data 3**).

To directly address whether each gene’s driver CpGs are organized around that gene’s own known regulators, we mapped each gene’s top extragenic driver CpGs to the nearest annotated gene within 500 kb and tested whether those neighboring genes share a known upstream transcription factor (TF) with the index gene, using TRRUSTv2 curated TF-target relationships and applying a hypergeometric test. Of the 45 testable genes (32 of 77 had no curated TRRUSTv2 regulator and were excluded), none reached nominal significance (smallest p = 0.71) (Supplementary **Fig. S3**, **Supplementary Data 4**). For *CD8A* specifically, whose driver-CpG neighborhoods we examined in detail, none of the 18 CpG-neighboring genes shared a RUNX3 regulatory relationship by either TRRUST curation or RUNX3 ChIP-seq peak overlap (0 of 18, versus a 1.3% genome-wide background rate). This gene-by-gene test reaches the same conclusion as the pooled analysis above: off-gene driver CpGs are not preferentially organized around their own target gene’s regulators.

### Extragenic CpGs are enriched within T cell enhancers but not at master-regulator binding sites

If predictive extragenic CpGs reflect lineage identity, they should be stably methylated and enriched within regulatory elements that establish and maintain T-cell fate. We therefore used Roadmap Epigenomics ChromHMM annotations and ChIP-seq peaks for lineage-defining transcription factors to determine whether predictive, extragenic CpGs localize to cis-regulatory elements or shared lineage transcription factor binding sites. The fraction of stable CpGs overlapping promoter-associated CpG islands was negatively associated with cross-validated R^2^ (r = −0.48; promoter CpG Islands are constitutively unmethylated in T cells and carry little inter-individual variance), while gene-body and open-sea CpGs dominated. High weighted predictive CpGs were predominantly annotated to T cell enhancer chromatin states in the Roadmap Epigenomics ChromHMM segmentation (enhancer fraction vs. cross-validated R2, r = +0.78; 23% of high-tier stable CpGs vs 12% for low-tier, **Fig. 2a, 2b**). This relationship was robust to the choice of ChromHMM reference: using PMA-ionomycin-stimulated Th/Th17 segmentations yielded a comparable association (r = +0.74), confirming that the enhancer enrichment pattern is not specific to the resting T cell DNA methylation state. Activation modestly increased enhancer annotation for some differentiation-associated genes (CCR6: +0.30, KLRG1: +0.09 enhancer fraction) but did not alter the overall performance gradient. The coupled CpGs avoided canonical, broadly active transcription-factor binding sites (*CTCF/SP1/ETS1* overlap vs cross-validated R^2^, r = −0.52, **Fig. 2c, 2d**).

**Figure 2.**
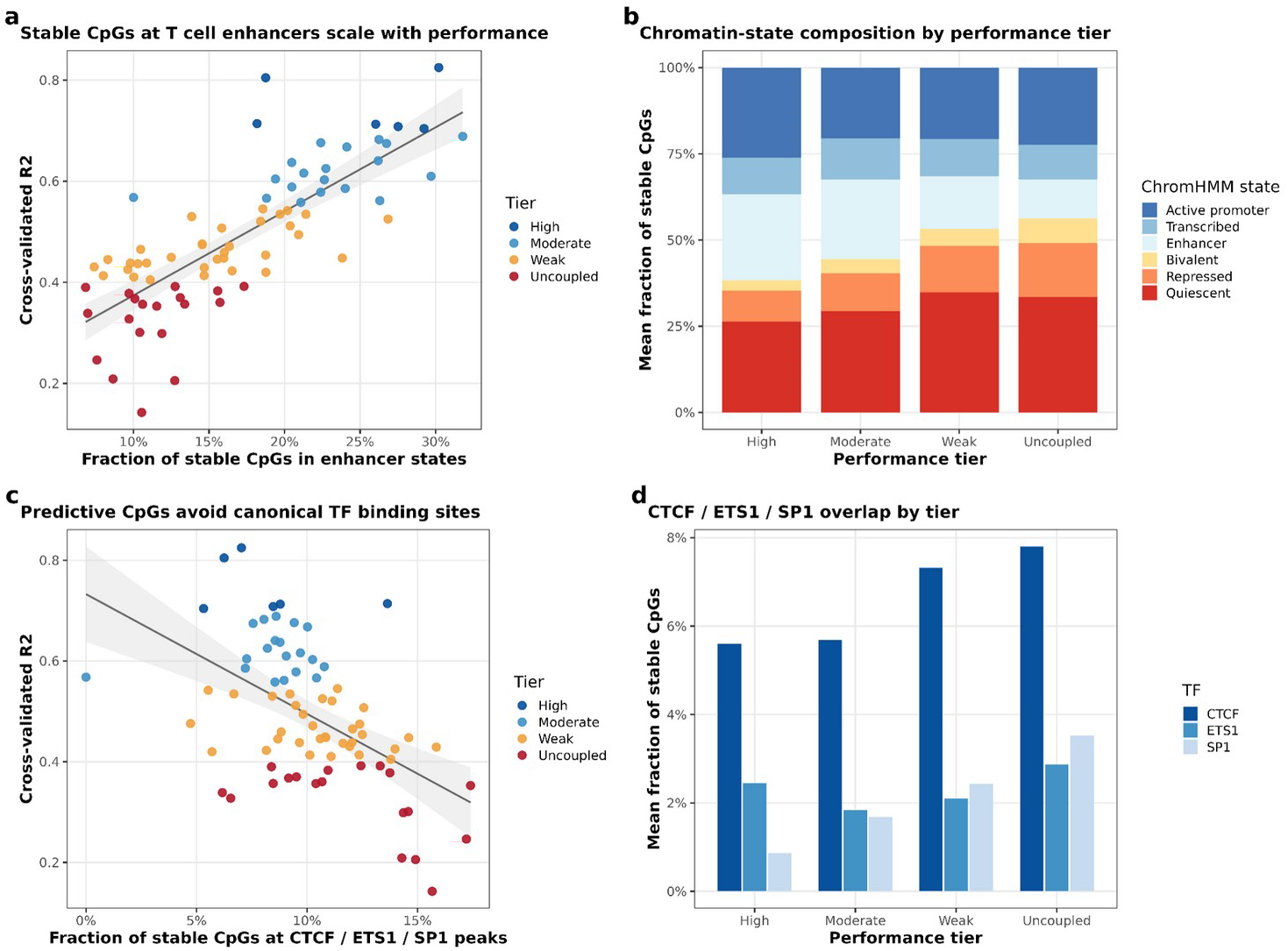
Off-gene CpGs are enriched within T cell enhancer chromatin states but are not at lineage master-regulator binding sites. **A)**, Fraction of a gene’s stable CpGs falling in Roadmap Epigenomics enhancer chromatin states (resting primary T cells: E043 CD4 naive, E044 CD4 memory, and E047 CD8 naive) versus cross-validated R^2^. Each point is one gene; color indicates performance tier; r = 0.78, n = 77 genes. **B)**, Mean chromatin-state composition of stable CpGs by performance tier, using the Roadmap Epigenomics 15-state ChromHMM core-marks model for resting primary T cells. Error bars show standard error across genes within each tier. The enhancer-state fraction is 23% for high-tier genes and 12% for low-tier genes. **C)**, Fraction of stable CpGs overlapping CTCF, ETS1, or SP1 ChIP-seq peaks (ENCODE) versus cross-validated R^2^. Broadly active canonical transcription-factor binding is negatively associated with model performance (r = −0.52, n = 77 genes). **D)**, Mean fraction of stable CpGs at CTCF, ETS1, and SP1 peaks by performance tier. High-tier genes show lower overlap with canonical transcription-factor sites than low-tier genes.

We next asked whether the extragenic CpGs are bound by lineage master regulators. Using primary Th1 T-bet/TBX21 ChIP-seq, and NK, plasma, and B cell ChIP-seq datasets for RUNX3, PRDM1, and FOXO1 (EOMES has no public immune ChIP-seq), we found that predictive extragenic CpGs were not enriched at master-regulator sites (**Fig. S4**). The apparent underrepresentation of extragenic CpGs at T-bet sites was likely an artifact of genomic context. Predictive CpGs are disproportionately located in open-sea regions, while T-bet occupancy is enriched at promoters and CpG islands. Accordingly, the signal vanished after stratification by CpG-island status (Cochran-Mantel-Haenszel (CMH) OR = 1.01, 95% CI = 0.99 to 1.03).

Notably, the top 10 predictive drivers for each gene remained de-enriched at T-bet sites with matched CpG context (island, shore, shelf, and open-sea) (CMH = OR 0.66, 95% CI = 0.48 to 0.91, p = 0.01). The host genes of the predictive extragenic CpGs were not associated with any specific biological program (0 GO terms at FDR < 0.05). Together, these results indicate that the predictive DNA methylation signal is likely distributed across enhancer DNA elements bound by factors other than the canonical lineage master regulators, T-bet, RUNX3, PRDM1, and FOXO1(**Fig. S4**).

### A lineage-versus-stimulus divide underlies gene predictability across multiple annotation databases

The nearly six-fold range in cross-validated R2 across the 77 T cell gene targets on this panel (0.14 to 0.82) could reflect genuine biology or a technical caveat of the panel; if biological, the well- and poorly-predicted genes should differ in function, not merely in how accurately they are measured. Enrichment across seven annotation databases drew a sharp line. Well-predicted genes were enriched for T cell differentiation and lineage commitment (GO Biological Process FDR = 1.3 x 10^-22, **Fig. 3a**; KEGG Th1/Th2/Th17 differentiation and T cell receptor signaling), whereas poorly predicted genes were enriched for negative regulators of T cell proliferation, immune checkpoints, and cytokines (FDR = 5.0 x 10^-14, **Fig. 3b**; KEGG inflammatory disease and FoxO signaling, the latter driven in part by FOXO1 itself, whose own expression is poorly predicted [CV R2 = 0.33] independent of its separate use above as a candidate regulatory input).

**Figure 3.**
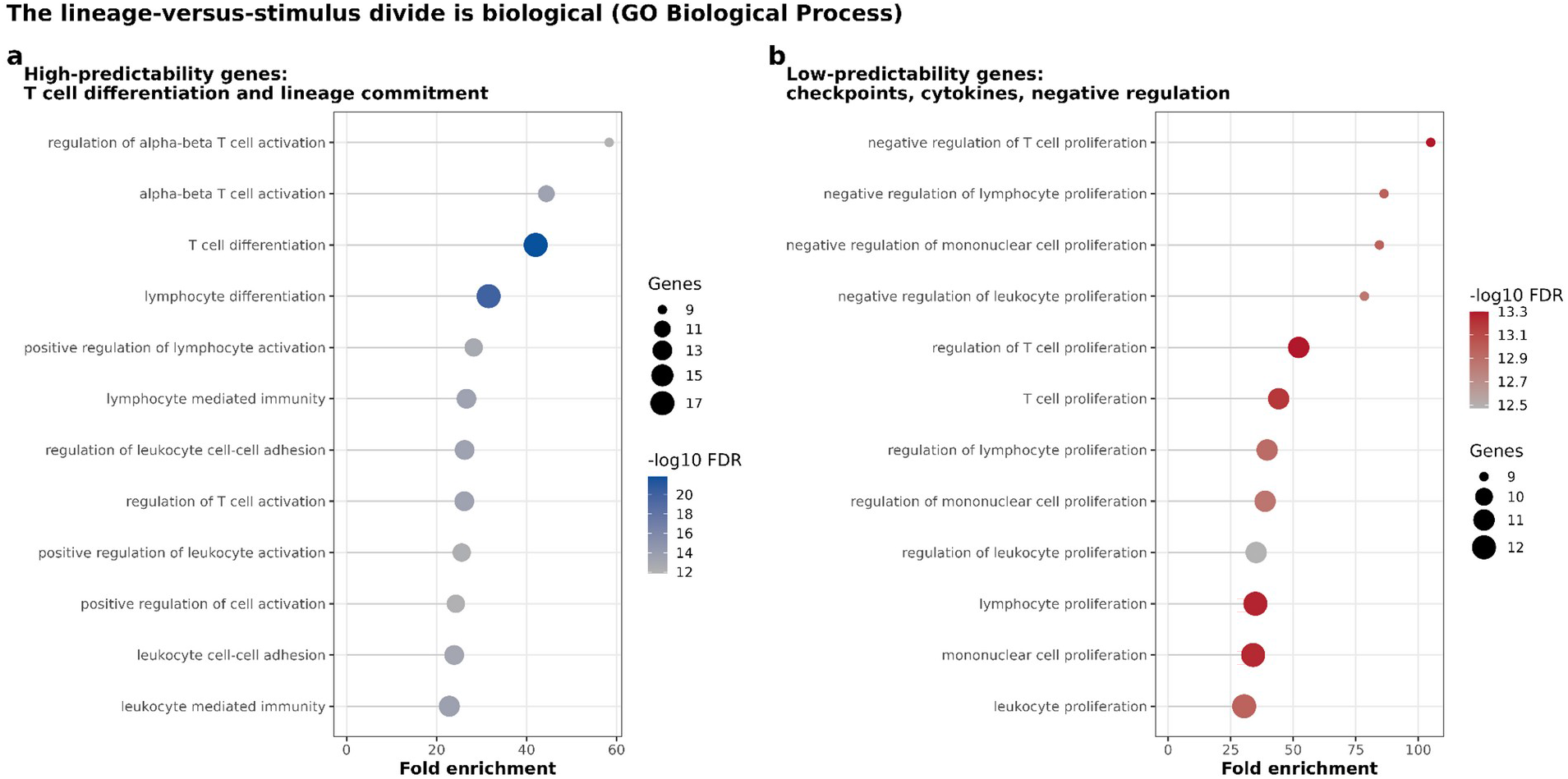
The lineage-versus-stimulus predictability divide is biological across annotation databases. **A)**, Gene Ontology Biological Process enrichment for the top tercile of genes by cross-validated R^2^ (high-predictability genes, n = 24). Dot size indicates the number of panel genes in each term; color indicates -log10 Benjamini-Hochberg FDR. The most significantly enriched term is “T cell differentiation” (FDR = 1.3 x 10^-22). **B)**, Gene Ontology Biological Process enrichment for the bottom tercile of genes by cross-validated R^2^ (low-predictability genes, n = 22). The most significantly enriched term is “negative regulation of T cell proliferation” (FDR = 5.0 x 10^-14). Enrichment was computed with clusterProfiler v4.14.6 against a genome-wide background. Fold enrichment on the x-axis is defined as the observed-to-expected gene ratio relative to the genome-wide background. Results across all seven annotation databases (Gene Ontology Biological Process, Molecular Function, Cellular Component, KEGG, Reactome, MSigDB Hallmark, and MSigDB C7 immunologic) are provided in **Supplementary Data S5**.

Molecular function enrichment analyses differentiated high-tier genes, characterized by DNA-binding transcription factor and coregulator activity, from low-tier genes, which were enriched for cytokine and kinase functions. The strongest discriminator was the MSigDB C7 immunologic collection, which identified 260 signatures associated with the high tier group (**Supplementary Data 5**). The expression of genes encoding transcription factors were predicted more accurately than non-transcription factors (TRRUST regulatory out-degree vs cross-validated R^2^, r = +0.29, **Fig. S5**). Consistent with the binding analysis above, this predictive signal was distributed across the methylome rather than concentrated at master-regulator binding sites.

### Methylation-predicted expression patterns validate in an independent cohort

Applied to an independent cancer cohort of 228 individuals (29 healthy donors, 106 patients with hematologic malignancies, and 93 patients with lung cancer), methylation-predicted expression remained highly concordant with measured expression for lineage and differentiation-associated genes (*CD3E* = 0.86, *CD3D* = 0.84, *CD28* = 0.83, *CD8A* = 0.82), whereas concordance was markedly lower for stimulus-responsive immune checkpoint ligands (*PD-L1* approximately 0.2, *PD-L2* approximately 0.1; panel-wide median cross-cohort Pearson r = 0.42), confirming the lineage-versus-stimulus divide in an independent cohort (**Fig. 4a**). Across donor diagnostic categories, methylation-predicted and measured expression remained highly concordant (group-mean correlations up to r = 0.97 for *CD8A*, 0.94 for *CD3E*), although variance was consistently compressed relative to measured expression values **(Fig. 4a, Supplemental Data 6)**. This pattern indicates that methylation-based estimates preserve biologically meaningful differences between groups while reducing within-group variation.

For the low-abundance senescence gene *CDKN2A/p16^INK4a^*, methylation-predicted expression distinguished diagnostic groups in an unadjusted, baseline-restricted comparison (Kruskal-Wallis χ^2^ = 25.5, df = 5, p = 1.1 × 10^-4^, n = 131), whereas measured NanoString expression did not (χ^2^ = 6.4, df = 5, p = 0.27, n = 108). In two-sided Wilcoxon tests versus healthy donors, predicted *p16^INK4a^* was nominally higher in plasma cell disorder (p = 2.3 × 10^-5^), chronic lymphocytic leukemia (p = 0.0018), and lung cancer (p = 0.0013), and did not differ significantly in lymphoma (p = 0.23) and acute leukemia (p = 0.90). The corresponding measured NanoString comparisons were nominally significant only for chronic lymphocytic leukemia (p = 0.049; all others p = 0.40 to 0.96).

Healthy donors in this cohort were substantially younger than every disease group (mean age 51.8 years versus 70 to 74 years for each malignancy), and predicted *p16^INK4a^*correlated with chronological age across the cohort (Pearson r = 0.385, p = 5.7 x 10^-6, n = 131 baseline samples; Methods). In the same baseline samples, *CD27* (r = −0.26, p = 0.0027), *CD28* (r = −0.23, p = 0.0098), and *CD127* (r = −0.23, p = 0.0080) each declined with age, though *CD8A* did not reach significance in this baseline cohort (r = −0.08, p = 0.36).

After age adjustment, none of the three nominal *p16^INK4a^* diagnosis effects remained significant (plasma cell disorder, p = 0.00055 to 0.11; chronic lymphocytic leukemia, p = 0.0044 to 0.089; lung cancer, p = 0.0031 to 0.47), and additional adjustment for cell composition did not restore significance (p = 0.56 to 0.92). The limited age overlap between healthy donors and cancer patients prevents a clear separation of age and diagnosis effects in this cross-sectional cohort. The unadjusted comparisons therefore do not establish disease-specific T-cell senescence. Predicted *p16^INK4a^* was associated with chronological age (Fig. 4c).

Because malignancies also shift blood composition, and because these samples are CD3-enriched but not FACS-pure (Methods), we re-tested every diagnosis-group difference in predicted expression for the full 8-gene panel (*CDKN2A/p16^INK4a^*, *CD8A*, *CD3E*, *GZMB*, *FOXP3*, *RUNX3*, *CD274*, *PDCD1*) with age and 12-cell composition adjustment together, restricted to one baseline (pre-treatment) sample per patient (n = 131; Methods). Calibration confirmed the expected CD3+ T-cell enrichment across most diagnoses, with chronic lymphocytic leukemia the exception, showing an elevated residual B cell fraction relative to every other group, consistent with circulating malignant B cells surviving the enrichment step in a disease where they dominate the lymphocyte pool (**Fig. 4b**). Of the 12 gene-diagnosis pairs nominally significant before adjustment, 4 remained significant after both age and composition adjustment together, all reflecting reduced predicted expression of lineage and checkpoint genes in acute leukemia relative to healthy donors: *CD8A* (p = 2.0 x 10^-4), *CD3E* (p = 1.4 x 10^-5), *RUNX3* (p = 1.9 x 10^-3), and *CD274*/PD-L1 (p = 1.2 x 10^-2). No *p16^INK4a^*association survived both adjustments, consistent with frequent CDKN2A/p16INK4A copy-number loss in acute leukemia rather than a regulatory, methylation-driven effect [14]. A diagnosis category present only in the pooled, multi-visit sample (“other hematologic malignancies”) does not correspond to any baseline patient population; it captured follow-up and on-treatment resamples with a missing diagnosis label and is not used in this baseline-restricted analysis (Methods). We report these four gene-diagnosis pairs as the composition- and age-robust disease associations in this cohort and do not extend the disease-specific interpretation further (**Fig. 4d; Supplementary Data 7**).

**Figure 4.**
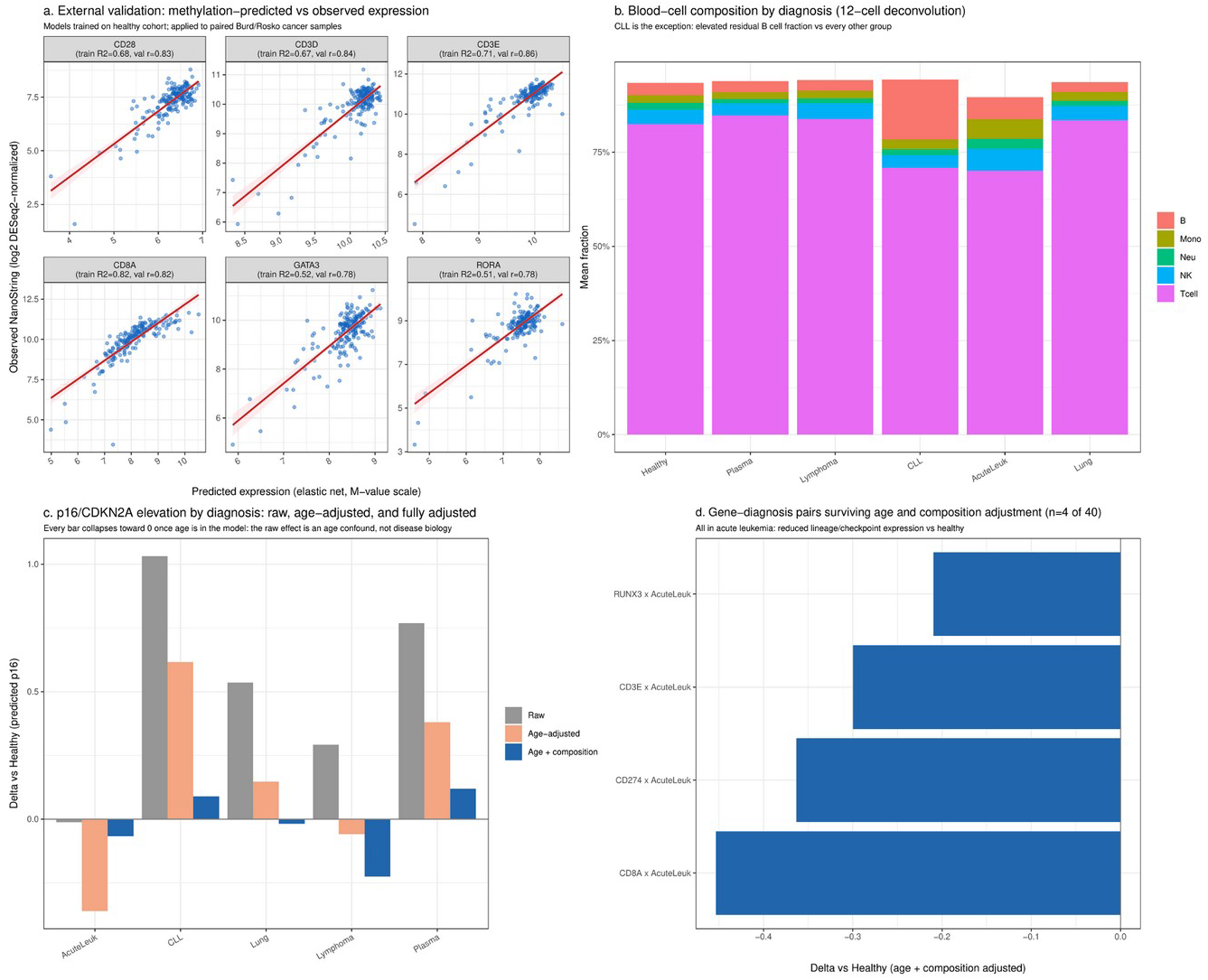
Methylation-predicted expression validates in an independent cancer cohort and recapitulates senescence biology. **A)** Scatterplots of methylation-predicted versus directly measured NanoString expression for six representative genes in the independent Burd/Rosko cancer cohort (n = 228; 29 healthy donors, 106 hematologic malignancies, and 93 lung cancers). Training cross-validated R^2^ and validation Pearson r are shown for each gene. The red line is a least-squares regression. Models were trained on the Ohio State MAG cohort and applied without retraining. **B)** Blood cell composition estimated by 12-cell EpiDISH deconvolution (cent12CT reference) by diagnosis group, restricted to one baseline sample per patient (n = 131). Chronic lymphocytic leukemia is the exception, with an elevated residual B cell fraction relative to every other group, consistent with circulating malignant B cells surviving the CD3-enrichment step. **C)** Methylation-predicted *p16/CDKN2A* elevation versus healthy donors by diagnosis group under three nested models: unadjusted (raw), age-adjusted, and age-plus composition-adjusted. Every bar collapses toward zero once age is included, indicating that the raw diagnosis effects are explained by the younger age of healthy donors (mean 51.8 years) relative to each malignancy group (mean 70 to 74 years) rather than disease-specific biology. **D)** Gene-diagnosis pairs surviving both age and composition adjustment (4 of 40 tested). All four reflect reduced predicted expression of lineage and checkpoint genes in acute leukemia relative to healthy donors. Full adjusted results are provided in **Supplementary Data S7.**

As a complementary test of the cell-fraction and detectability logic, we asked whether methylation-expression coupling strengthens for a gene in the disease subgroup where it is most highly expressed, relative to healthy donors. We identified 15 candidate genes with weak overall predictability (cross-validated R^2^ < 0.45) that were nonetheless substantially elevated in at least one disease group relative to healthy donors (delta > 0.5 log2 units; for example *IL6* in plasma cell disorder, *NECTIN2* and *CDKN2A//p16^INK4a^* in acute leukemia, and *CDKN2A/ARF* and *CD276* in other hematologic malignancies). For each gene, we computed the mean absolute correlation between methylation at its top 50 highest-weight model CpGs and measured expression, separately within each adequately powered diagnosis group (n >= 14: acute leukemia, healthy donors, other hematologic malignancies, lung cancer, and plasma cell disorder; chronic lymphocytic leukemia and lymphoma were underpowered and excluded), and compared each gene’s coupling in healthy donors to its coupling in the disease group where its expression was highest. Coupling strengthened in the higher-expression group for 12 of 15 genes (binomial p = 0.035), with the largest gains for *PVR* (mean absolute r: 0.138 in healthy donors to 0.277 in acute leukemia), *NECTIN2* (0.130 to 0.251 in acute leukemia), and *CDKN2A/ARF* (0.164 to 0.263 in other hematologic malignancies). The three exceptions were *CEACAM1* (a marginal decline, 0.184 to 0.178, in other hematologic malignancies), *BTLA* (no disease group exceeded healthy expression, precluding a directional test), and *PDCD1LG2* (a decline, 0.167 to 0.098, in lung cancer). This pattern is consistent with the mechanism proposed throughout: weak predictability in the training cohort partly reflects insufficient expression variance and cell-fraction dilution rather than the complete absence of an underlying methylation-expression relationship, which becomes detectable once expression rises in a relevant disease context (**Supplementary Data 8**).

### The lineage signal is a stable genomic property robust to cellular admixture

T cell lineage genes are expressed and epigenetically regulated in a cell-type-specific manner; if the methylation signal underlying their prediction were merely a proxy for T cell abundance rather than a stable genomic feature, it would be expected to degrade or disappear when applied to PBMC samples containing a mixture of T, B, NK, and monocyte cells. To test this directly, we applied the T cell models to matched whole-PBMC methylation from 385 individuals from the same cohort. By deconvolution, these samples contained a median T cell fraction of 50%. Methylation-expression models for lineage and differentiation-associated genes retained performance despite the admixture (*CD8A* r = 0.86, matching its T cell test performance; *KLRG1* = 0.79, *GZMB* = 0.77, *TBX21* = 0.80, *EOMES* = 0.72, **Fig. 5a**; **Fig. S6**). Critically, the observed signal was not driven by cell composition: adjusting for T cell fraction left the median correlation essentially unchanged (0.534 to 0.543), and adjusting for the full 12-cell composition reduced it only modestly (to 0.514), with the high-weight lineage genes surviving full adjustment (*CD8A* = 0.72, *KLRG1* = 0.71, *GZMB* = 0.66, *TBX21* = 0.62). Two genes were exceptions and collapsed under adjustment for composition: *SPI1*, because *PU.1/SPI1* is expressed in B and myeloid cells, and *CD3E*, a pan-T marker whose methylation tracks the T cell fraction rather than a stable lineage program (**Fig. S7**). Although PBMC-trained models predicted lineage- and differentiation-associated genes with similar accuracy, they shared only ∼1.5% of CpGs with the T-cell models (**Fig. 5b,c; Supplementary Data 9**), indicating that the same lineage signal can be encoded by many alternative CpG combinations. The predictive methylation pattern at lineage genes is therefore a stable genomic feature.

**Figure 5.**
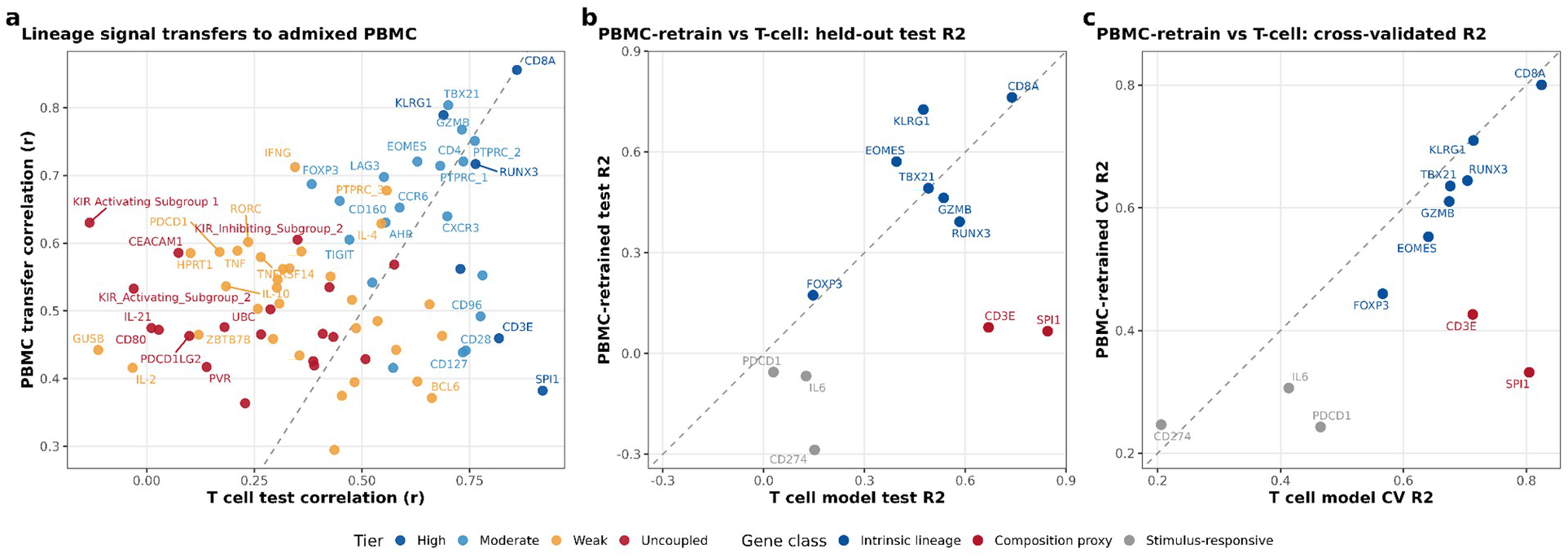
The lineage signal is a stable genomic property robust to cellular admixture. **A)**, Transfer of T cell-trained models to whole-PBMC methylation. Each point is one gene, showing the Pearson correlation between methylation-predicted expression (from PBMC methylation) and directly measured purified-T cell NanoString expression, plotted against the T cell model’s held-out test correlation. Color indicates performance tier; gene class (intrinsic lineage, composition proxy, stimulus-responsive) is indicated by point shape. Genes above the identity line (dashed) transfer comparably or better from PBMC; most intrinsic lineage genes cluster near the diagonal. Labeled outliers SPI1 and CD3E are composition-driven (see text). n = 362 matched PBMC-and-T cell sample pairs. **B)**, Held-out test R^2^ for models retrained directly on PBMC methylation (y-axis) versus the original T cell models (x-axis). Each point is one gene. Robust lineage genes (*CD8A*, *KLRG1*, *EOMES*, and *GZMB*) retrain comparably from PBMC; composition-driven genes (*SPI1* and *CD3E*, red labels) and stimulus-responsive genes perform poorly under both model sets. **C)**, Same comparison using cross-validated R^2^ in place of held-out test R^2^, confirming the held-out result is not an overfitting artifact. The dashed line represents identity in all panels. n = 385 PBMC samples (training for PBMC-retrained models); 20% held-out test set for panel b.

### Methylation-predicted scores generalize to whole blood at population scale

To test whether the lineage signal generalizes beyond the training context, we applied the PBMC-weighted models to the MGB/Harvard whole-blood biobank (n = 4,483 with age data; n = 4,250 with complete outcome follow-up, median 8.6 years). As a validity check, predicted *p16/CDKN2A* increased with chronological age after full composition adjustment (beta = +0.020 SD per year, p = 5.0 x 10^-105), and naive T cell lineage scores (*CD27*, *CD28*, *CD8A, CD127*, *CD3E*) declined as expected, confirming the methylation signal generalizes across cohorts, sample types, and independently performed normalization pipelines (**Fig. S8**). Exploratory associations with incident disease and all-cause mortality are reported in the Supplementary Information; the strongest signal was a composition-adjusted association between predicted *CDKN2A/ARF* expression and incident COPD, consistent across both T cell and PBMC model sets (**Fig. S9**; **Supplementary Data 10-11**). One additional nominal association — between the T cell-weighted CD80 score and incident asthma (OR = 1.24, BH-FDR = 0.030; Fig. S9a) — is not interpreted as reflecting CD80 biology, given that this gene’s model showed negative held-out predictive validity in the primary T cell cohort (Supplementary Data 2).

## Discussion

Prediction accuracy varied with gene function and the regulatory annotations of predictive CpGs. Genes defining stable lineage and differentiation programs were predicted with high accuracy, whereas stimulus-responsive genes, including immune checkpoints and cytokines, were consistently poorly predicted despite comparable expression levels. This distinction was evident before model fitting, as low-performing genes showed weak methylation-expression coupling at individual CpGs, indicating that the observed performance gradient reflects fundamental differences in regulatory mode rather than measurement sensitivity or insufficient model complexity (Fig. 1a, b; Fig. S1).

The predictive signal was encoded predominantly outside annotated gene bodies and was strongly concentrated in distal enhancer elements. Restricting models to gene-associated CpGs substantially reduced predictive performance, often abolishing signal altogether, demonstrating that extragenic CpGs contribute essential regulatory information rather than redundant correlates. Genome-wide, high-weight driver CpGs were enriched within active T-cell enhancers, and enhancer representation strongly tracked model performance. Notably, this association was preserved when enhancer states were defined in activated T cells, indicating that predictive information resides within stable lineage-associated regulatory architecture rather than transient activation-induced elements. These findings are consistent with evidence that lineage-specific enhancer demethylation is established during T-cell differentiation and maintained as a durable epigenetic feature of cell identity [11, 15, 16]. They also align with eQTM studies showing that methylation-expression relationships are preferentially enriched at distal regulatory elements rather than promoters, suggesting that enhancer methylation constitutes a major conduit through which DNA methylation captures transcriptional state (Fig. 1b, 2a, b)[7, 8].

Predictive CpGs were overwhelmingly gene-specific, showed little recurrence across models, and failed to converge on canonical lineage master-regulator binding sites or coherent upstream pathways. Although T-cell differentiation is orchestrated by factors such as TBX21, EOMES, and related lineage regulators, our results indicate that their downstream transcriptional outputs are recorded by methylation marks distributed across a broad enhancer landscape rather than through a small number of shared regulatory loci. This observation is consistent with epigenomic models in which cell identity is maintained through large networks of lineage-associated enhancers whose collective activity is highly redundant and robust to perturbation of individual regulatory elements [17, 18]. This distributed pattern was also evident at the transcription-factor-binding level: driver CpGs showed no enrichment at canonical lineage master-regulator binding sites in the assessed ChIP-seq datasets (Fig. 2c, d; Fig. S4, S5). Consistent with this distributed, non-centralized regulatory logic, transcription-factor target gene sets have separately been shown to support an explainable methylation-based predictor of chronological age [19], in which 10,000 age-correlated CpGs were linked through 2,421 annotated genes to 1,137 transcription-factor target gene sets.

The mechanistic distinction between predictable and unpredictable genes reflects a broader division between constitutive and inducible transcriptional programs. Lineage-defining genes acquire expression states during differentiation and maintain them through stable cis-regulatory chromatin architecture, generating an enduring molecular record that remains detectable in circulating lymphocytes. In contrast, genes such as *CD274*, *PDCD1LG2*, *IL6*, and *TNFA* are controlled by rapidly responsive signaling pathways, including JAK-STAT-, IRF1-, and NF-κB-mediated transcriptional circuits, and are additionally influenced by post-transcriptional mechanisms such as RNA-binding proteins and AU-rich-element-dependent mRNA turnover [12, 13, 20]. These pathways regulate transient cellular responses rather than differentiation history, providing a biological explanation for their limited predictability from methylation.

This interpretation was reinforced in an independent validation cohort. Methylation-predicted expression closely tracked measured expression for stable lineage markers but not for stimulus-responsive genes, demonstrating that the models capture reproducible biological signal rather than overfitting to the training dataset (Fig. 4a). Moreover, methylation-derived *CDKN2A*/*p16^INK4a^* scores distinguished diagnosis groups in an unadjusted comparison more effectively than directly measured transcript abundance, an effect fully explained by the age difference between healthy donors and disease groups rather than disease-specific senescence biology (Fig. 4c). This finding is consistent with prior work establishing *p16^INK4a^* as a robust marker of T-cell aging in vivo [21, 22] and suggests that methylation-derived proxies may, in some contexts, provide a more stable readout of long-term cellular state than direct transcript measurements, which are often influenced by stochastic and short-term fluctuations.

The transferability of the signal from sorted T cells to PBMC and whole-blood datasets further supports this interpretation. Predictive performance remained largely intact following adjustment for inferred cellular composition, and PBMC-trained models reproduced comparable accuracy despite using largely non-overlapping CpG sets. Together, these findings suggest that lineage-associated methylation information is encoded redundantly across many genomic loci and can therefore be recovered through multiple alternative CpG combinations. Such redundancy may explain why methylation-derived biomarkers often generalize successfully across cohorts despite limited overlap in individual CpG features (Fig. 5a-c; Supplementary Data S9).

These findings have implications beyond T-cell biology. Many widely used methylation-derived surrogates, including GrimAge, GrimAge2, DNAmFitAge, OMICmAge, and methylation-based protein EpiScores, are selected primarily for predictive performance, while the regulatory basis of their signal remains largely unknown. Our results suggest that the achievable performance and transferability of any methylation proxy are mechanistically bounded by the regulatory architecture of the target phenotype. Traits encoded through stable enhancer-associated differentiation programs should be more amenable to methylation-based prediction than phenotypes governed by acute signaling, post-transcriptional regulation, or multi-step biochemical cascades. More broadly, the enhancer enrichment analyses and on-gene retraining controls described here provide a general framework for distinguishing mechanistically grounded methylation proxies from predictors that rely primarily on indirect statistical associations.

In summary, blood DNA methylation encodes T-cell lineage identity through distributed enhancer-centered regulatory programs that are stable across activation states, reproducible across cohorts, and transferable across biospecimens. These findings define an important boundary for methylation-based molecular prediction: DNA methylation is well suited to capturing durable differentiation history but is inherently less effective for reconstructing transient stimulus-responsive transcriptional states.

## Supporting information

Supplementary Data Legends for S1 - S11

## Acknowledgments

We thank Kristen Seale, Natalia Carreras-Gallo, Laura Balague, and Mike Mallin of the TruDiagnostic Research and Bioinformatics teams for the discussion that inspired this analysis and shaped the current form of the manuscript. We also thank Janice K. Kiecolt-Glaser for her contribution of the cancer-cohort samples used in this study. We also thank Adiv A. Johnson for his review of and edits to the manuscript.

## Funding

No funding was received for this study.

## Author contributions

- V.B.D.: Conceptualization, methodology, formal analysis, software, and writing of the original draft
- M.M.: Formal analysis, review and editing
- S.A.H.: Methodology, review and editing
- A.A.: Review and editing
- A.E.R: Cancer-cohort samples and clinical-data generation
- C.P.: Training-cohort sample and data generation
- R.S.: Resources, review and editing
- J.L.S: Validation dataset
- C.E.B.: supervision, resources, writing (review and editing)

## Disclosure of interest

V.B.D., S.A.H., and R.S. are employees of TruDiagnostic Inc., which commercializes DNA methylation testing. J.L.S. is an advisor for TruDiagnostic Inc. The remaining authors declare no competing interests.

## Declaration of generative AI use

Generative AI (OpenAI Codex, OpenAI) was used for formatting checks, error identification, and limited language editing. Generative AI was not used to perform scientific analyses.

## Data availability statement

Raw methylation IDAT files, raw NanoString expression data, and normalized beta matrices for the Ohio State MAG training cohort, the PBMC cohort, and the Burd/Rosko cancer cohort cannot be made publicly available due to institutional data-access agreements and participant privacy restrictions. Per-sample methylation-predicted expression scores (a derived, not raw, quantity) for the cancer cohort (N = 228) are provided as Supplementary Data S6, together with the age- and composition-adjusted disease-association results (Supplementary Data S7), the coupling-shift per-gene results (Supplementary Data S8), and Supplementary Data S1 to S5 and S9 to S11 (covering the remaining processed/derived results referenced throughout this manuscript). All eleven Supplementary Data files (S1 to S11), the complete Supplementary Information document (including supplementary methods, notes, figures, and embedded tables), and the Supplementary Data legends are publicly available in Zenodo at https://doi.org/10.5281/zenodo.23185649. The same repository record provides the analysis code and figure files. The Mass General Brigham Biobank data are available to qualified investigators through the MGB Biobank data-access process (https://biobank.partners.org); the disease-outcome modeling for that cohort was performed by the MGB/Harvard collaborators against their institutional clinical database and is not independently reproducible from data available to the corresponding authors (see Code availability). Chromatin state maps, TF ChIP-seq peaks, pathway databases, and all other external reference resources are publicly available; accessions, versions, and download URLs are listed in Supplementary Note 1 (deposited with the Zenodo code archive as DATA_SOURCES.md; this is a documentation file, not one of the numbered Supplementary Data tables, and is labeled as a Note specifically to avoid collision with the Supplementary Data S1 to S11 numbering).

## Code availability statement

Analysis code for all figures, tables, and statistical analyses reported in this study, together with the processed/derived Supplementary Data tables (S1 to S11) and the main and supplementary figures, is available at https://doi.org/10.5281/zenodo.23185649. The archive does not include the trained methylation-to-expression models or any per-CpG model coefficient table: the 77 gene-specific elastic nets (T cell and PBMC-admixture-aware sets) plus the portable deployment bundle are available from the corresponding author (V.B.D.) after signing a Data Use Agreement, because the training data cannot be shared under the applicable data-access agreements and per-CpG model coefficients would allow the models to be reconstructed from otherwise-shareable result tables. The Zenodo archive’s own README documents this exclusion, and every other processed result needed to verify the manuscript’s statistical claims, in full.

## Notes

https://doi.org/10.5281/zenodo.23185649

### Summary of Updates

This version has been revised to improve clarity and presentation. Repetitive interpretive statements and formulaic transitions have been reduced, including a shorter conclusion to the abstract. The p16 results have been reorganized to distinguish unadjusted diagnosis comparisons, age associations, and adjusted analyses. The text now emphasizes that limited age overlap prevents a clear separation of age and diagnosis effects, and that the unadjusted comparisons do not establish disease-specific T-cell senescence. Mathematical notation, superscripts, spacing, and manuscript layout have been corrected. The Methods section has been moved before the Results, and author contact information and declarations have been updated. The Data and Code Availability statements now cite the updated Zenodo deposit, DOI 10.5281/zenodo.23185649, which includes the complete supplementary information, data legends, analysis code, figures, and all eleven supplementary data files. The underlying analyses and reported numerical results have not been changed.

## References

1. Lu, A. T. et al. DNA methylation GrimAge strongly predicts lifespan and healthspan. Aging (Albany NY) 11, 303–327 (2019).

2. Lu, A. T. et al. DNA methylation GrimAge version 2. Aging (Albany NY) 14, 9484–9549 (2022).

3. Gadd, D. A. et al. Epigenetic scores for the circulating proteome as tools for disease prediction. eLife 11, e71802 (2022).

4. McGreevy, K. M. et al. DNAmFitAge: biological age indicator incorporating physical fitness. Aging (Albany NY) 15, 3904–3938 (2023).

5. Arpawong, T. E. et al. Physiological health Age (PhysAge): a novel multi-system molecular timepiece predicts health and mortality in older adults. GeroScience 48, 3115–3135 (2026).

6. Chen, Q. et al. OMICmAge quantifies biological age by integrating multi-omics with electronic medical records. *Nat*. Aging 6, 722–737 (2026).

7. Kim, S. et al. Cis- and trans-eQTM analysis reveals novel epigenetic and transcriptomic immune markers of atopic asthma in airway epithelium. J. Allergy Clin. Immunol. 152, 887–898 (2023).

8. Keshawarz, A. et al. Expression quantitative trait methylation analysis elucidates gene regulatory effects of DNA methylation: the Framingham Heart Study. Sci. Rep. 13, 12952 (2023).

9. Harland, K. L. et al. Epigenetic plasticity of the Cd8a locus during CD8+ T-cell development and effector differentiation and reprogramming. Nat. Commun. 5, 3547 (2014).

10. Cruz-Guilloty, F. et al. Runx3 and T-box proteins cooperate to establish the transcriptional program of effector CTLs. J. Exp. Med. 206, 51–59 (2009).

11. Nestor, C. E. et al. 5-Hydroxymethylcytosine remodeling precedes lineage specification during differentiation of human CD4+ T cells. Cell Rep. 16, 559–570 (2016).

12. Matsusaka, T., et al. Transcription factors NF-IL6 and NF-kappa B synergistically activate transcription of the inflammatory cytokines, interleukin 6 and interleukin 8. Proc. Natl Acad. Sci. USA 90, 10193–10197 (1993).

13. Garcia-Diaz, A. et al. Interferon receptor signaling pathways regulating PD-L1 and PD-L2 expression. Cell Rep. 19, 1189–1201 (2017).

14. Cayuela, J. M., Hebert, J. & Sigaux, F. Homozygous MTS1 (p16INK4A) deletion in primary tumor cells of 163 leukemic patients. Blood 85, 854 (1995).

15. Lal, G. et al. Epigenetic regulation of Foxp3 expression in regulatory T cells by DNA methylation. J. Immunol. 182, 259–273 (2009).

16. Toker, A. et al. Active demethylation of the Foxp3 locus leads to the generation of stable regulatory T cells within the thymus. J. Immunol. 190, 3180–3188 (2013).

17. Hnisz, D. et al. Super-Enhancers in the Control of Cell Identity and Disease. Cell 155, 934–947 (2013).

18. Whyte, W. A. et al. Master transcription factors and mediator establish super-enhancers at key cell identity genes. Cell 153, 307–319 (2013).

19. Shokhirev, M. N. & Johnson, A. A. Using buccal methylomic data to create explainable aging clocks as well as classifiers and regressors for lifestyle and demographic factors. Front. Genet. 16, 1637186 (2025).

20. Guo, L., Vlasova-St Louis, I. & Bohjanen, P. R. Post-transcriptional regulation of cytokine expression and signaling. Curr. Trends Immunol. 19, 33–40 (2018).

21. Liu, Y. et al. Expression of p16INK4a in peripheral blood T-cells is a biomarker of human aging. Aging Cell 8, 439–448 (2009).

22. Nelson, J. A. E. et al. Expression of p16INK4a as a biomarker of T-cell aging in HIV-infected patients prior to and during antiretroviral therapy. Aging Cell 11, 916–918 (2012).

23. Aryee, M. J. et al. Minfi: a flexible and comprehensive Bioconductor package for the analysis of Infinium DNA methylation microarrays. Bioinformatics 30, 1363–1369 (2014).

24. Fortin, J.-P. et al. Functional normalization of 450k methylation array data improves replication in large cancer studies. Genome Biol. 15, 503 (2014).

25. Niu, L., Xu, Z. & Taylor, J. A. RCP: a novel probe design bias correction method for Illumina Methylation BeadChip. Bioinformatics 32, 2659–2663 (2016).

26. Triche, T. J. Jr, Weisenberger, D. J., Van Den Berg, D., Laird, P. W. & Siegmund, K. D. Low-level processing of Illumina Infinium DNA Methylation BeadArrays. Nucleic Acids Res. 41, e90 (2013).

27. Risso, D., Ngai, J., Speed, T. P. & Dudoit, S. Normalization of RNA-seq data using factor analysis of control genes or samples. Nat. Biotechnol. 32, 896–902 (2014).

28. Love, M. I., Huber, W. & Anders, S. Moderated estimation of fold change and dispersion for RNA-seq data with DESeq2. Genome Biol. 15, 550 (2014).

29. Ritchie, M. E. et al. limma powers differential expression analyses for RNA-sequencing and microarray studies. Nucleic Acids Res. 43, e47 (2015).

30. Smyth, G. K. Linear models and empirical Bayes methods for assessing differential expression in microarray experiments. Stat. Appl. Genet. Mol. Biol. 3, Article 3 (2004).

31. Friedman, J., Hastie, T. & Tibshirani, R. Regularization paths for generalized linear models via coordinate descent. J. Stat. Softw. 33, 1–22 (2010).

32. Han, H. et al. TRRUST v2: an expanded reference database of human and mouse transcriptional regulatory interactions. Nucleic Acids Res. 46, D380–D386 (2018).

33. Roadmap Epigenomics Consortium et al. Integrative analysis of 111 reference human epigenomes. Nature 518, 317–330 (2015).

34. Lawrence, M. et al. Software for computing and annotating genomic ranges. PLoS Comput. Biol. 9, e1003118 (2013).

35. Oki, S. et al. ChIP-Atlas: a data-mining suite powered by full integration of public ChIP-seq data. EMBO Rep. 19, e46255 (2018).

36. Hammal, F., de Langen, P., Bergon, A., Lopez, F. & Ballester, B. ReMap 2022: a database of human, mouse, Drosophila and Arabidopsis regulatory regions from an integrative analysis of DNA-binding sequencing experiments. Nucleic Acids Res. 50, D316–D325 (2022).

37. Yu, G. & He, Q.-Y. ReactomePA: an R/Bioconductor package for Reactome pathway analysis and visualization. Mol. BioSyst. 12, 477–479 (2016).

38. Wu, T. et al. clusterProfiler 4.0: a universal enrichment tool for interpreting omics data. Innovation (Camb.) 2, 100141 (2021).

39. Phipson, B., Maksimovic, J. & Oshlack, A. missMethyl: an R package for analyzing data from Illumina’s HumanMethylation450 platform. Bioinformatics 32, 286–288 (2016).

40. Teschendorff, A. E., Breeze, C. E., Zheng, S. C. & Beck, S. A comparison of reference-based algorithms for correcting cell-type heterogeneity in Epigenome-Wide Association Studies. BMC Bioinformatics 18, 105 (2017).

41. Houseman, E. A. et al. DNA methylation arrays as surrogate measures of cell mixture distribution. BMC Bioinformatics 13, 86 (2012).

42. Salas, L. A. et al. Enhanced cell deconvolution of peripheral blood using DNA methylation for high-resolution immune profiling. Nat. Commun. 13, 761 (2022).

43. Luo, Q. et al. A meta-analysis of immune-cell fractions at high resolution reveals novel associations with common phenotypes and health outcomes. Genome Med. 15, 59 (2023).

44. Therneau, T. M. & Grambsch, P. M. Modeling Survival Data: Extending the Cox Model (Springer, New York, 2000).

45. Therneau, T. M. A Package for Survival Analysis in R. R package version 3.7–0 (2024).

