## Supplementary Data Legends for S1 - S11 for "Regulatory mode and enhancer architecture determine limits of blood DNA methylation as a molecular proxy"

**Supplementary Data S1. Pre-amplified panel gene performance and expression abundance.** Per-gene methylation-to-expression model performance (cross-validated $R^2$ and held-out test $R^2$) and mean log2-normalized NanoString expression for all 77 T cell genes, annotated by whether the gene was measured using pre-amplification chemistry (pre-amplified call set: *CDKN2A/p16*, *CDKN2A/ARF*, *IL-17A*, *IL-6*, *FOXO1*, and *CDKN1B*) or standard NanoString chemistry. Pre-amplified genes had lower median cross-validated $R^2$ (0.34 vs 0.48, Wilcoxon $p = 0.0017$) but were not lower in mean expression abundance (Wilcoxon $p = 0.30$), indicating their weaker coupling reflects stimulus-responsive biology rather than a detection-floor artifact.

**Supplementary Data S2. On-gene-only versus genome-wide model performance for all 77 T cell genes.** Per-gene comparison of held-out test $R^2$ for elastic net models restricted to each gene's own annotated CpGs (on-gene-only) versus the standard genome-wide top-50,000 CpG model, evaluated on the same subject-level 80/20 held-out split. Columns include: gene, cross-validated $R^2$ of the genome-wide model, number of on-target CpGs available, on-gene-only held-out test $R^2$, genome-wide held-out test $R^2$, performance tier, and delta (genome-wide minus on-gene-only test $R^2$). Genes with NA for on-gene-only test $R^2$ had zero stable CpGs annotated to the target gene. Negative on-gene-only test $R^2$ values indicate models that performed worse than predicting the mean, confirming that on-gene CpGs alone carry insufficient signal for many genes.

**Supplementary Data S3. Per-gene on-gene versus off-gene CpG audit for all 77 T cell elastic net models.** For each of the 77 T cell genes, the table reports the cross-validated $R^2$ and performance tier; the total number of stable CpGs selected by the elastic net (present in at least 2 of 3 cross-validation folds); and the breakdown of those CpGs by genomic location relative to the target gene: on-gene (annotated to the target gene's own promoter or body), other-gene (annotated to a different gene's promoter or body), or distal/intergenic. For each location category, both the CpG count share (percentage of total stable CpGs) and the model weight share (sum of absolute elastic-net coefficients for that category divided by total) are reported, along with the maximum absolute elastic-net coefficient for on-gene and other-gene CpGs respectively. Genes are ordered by cross-validated $R^2$ (descending).

**Supplementary Data S4. Gene-by-gene transcription factor module enrichment analysis for all 77 T cell genes.** For each of the 77 T cell genes, the table reports the performance tier and cross-validated $R^2$; whether the gene was highlighted as a named worked example; the number of top off-gene driver CpGs tested (top 20 by absolute elastic-net weight among stable CpGs) and the number of unique neighboring genes identified within 500 kb (hg19). The TRRUST v2 analysis columns report the number and identity of curated upstream transcription factor regulators for the index gene, whether the gene was testable (at least one curated TRRUST regulator), the number and identity of neighboring genes sharing at least one upstream regulator with the index gene, and the hypergeometric p-value (population = all TRRUST-annotated genes; genes with no curated TRRUST regulator are marked not testable and assigned no p-value). The ChIP-seq sensitivity columns report the best-performing lineage transcription factor (TBX21, RUNX3, FOXO1, or PRDM1) for each gene's neighboring-gene promoter overlap, the cell-type context of that ChIP-seq dataset, whether the index gene's own promoter is bound, the number and fraction of neighboring gene promoters bound, the genome-wide background promoter-binding rate for that transcription factor, and the hypergeometric p-value.

**Supplementary Data S5. Functional enrichment results across seven annotation databases.** Results of tier-stratified functional enrichment comparing the top and bottom terciles of genes by cross-validated $R^2$ (high-predictability n = 24 genes; low-predictability n = 22 genes) across seven annotation databases: Gene Ontology Biological Process, Molecular Function, and Cellular Component; KEGG; Reactome; MSigDB Hallmark; and MSigDB C7 immunologic gene sets. Enrichment was computed with clusterProfiler v4.14.6 (Gene Ontology and KEGG), ReactomePA v1.50.0 (Reactome), and msigdbr v25.1.1 (MSigDB). P-values were corrected by Benjamini-Hochberg FDR within each collection. Columns include: collection, tier, term description, FDR-adjusted p-value, gene count, and gene identifiers.

**Supplementary Data S6. Methylation-predicted expression scores for all 77 genes in the independent cancer validation cohort.** Per-sample methylation-predicted expression scores for all 77 T cell genes in the Burd/Rosko cancer cohort (n = 228; 29 healthy donors, 106 hematologic malignancies, 93 lung cancers). Scores are on the M-value scale (logit2-transformed beta). Columns include sample identifier, clinic, diagnosis group, age, sex, and one column per predicted gene. These data underlie the validation results in Fig. 4 and can be used to reproduce the cross-cohort concordance, p16 diagnosis comparisons, and composition-adjustment analyses.

**Supplementary Data S7. Composition- and age-adjusted disease-diagnosis associations with predicted expression in the independent cancer cohort.** Linear-model results testing the association between diagnosis group (versus healthy donors) and methylation-predicted expression for 8 T cell genes (*CDKN2A/p16*, *CD8A*, *CD3E*, *GZMB*, *FOXP3*, *RUNX3*, *CD274*, *PDCD1*) across 5 diagnosis groups (plasma cell disorder, lymphoma, chronic lymphocytic leukemia, acute leukemia, lung cancer) in the Burd/Rosko cancer cohort, restricted to one baseline (pre-treatment) sample per patient (n = 131 with complete chronological age data). A sixth, pooled-sample category ("other hematologic malignancies") reflected follow-up/on-treatment resamples with a missing diagnosis label and is excluded here. Each gene-diagnosis pair is tested under three nested linear models: unadjusted (diagnosis only), age-adjusted (diagnosis plus chronological age), and fully adjusted (diagnosis, age, and 12-cell EpiDISH composition). Columns: gene; group (diagnosis group); n; raw_delta and raw_p (unadjusted model coefficient and p-value); age_delta and age_p (age-adjusted model); full_delta and full_p (age- and composition-adjusted model); survives_age and survives_full (TRUE if $p < 0.05$ with effect in the same direction as the unadjusted model). These data underlie Fig. 4d and show that most raw diagnosis effects, including all *CDKN2A/p16* associations, are attributable to the age difference between healthy donors and cancer patients in this cohort (mean age 51.8 versus 70 to 74 years across malignancy groups) rather than disease-specific biology; four gene-diagnosis pairs, all in acute leukemia, remain significant after both age and composition adjustment.

**Supplementary Data S8. Methylation-expression coupling shift by disease-expression level for 15 weakly predictable candidate genes in the cancer cohort.** For each of 15 T cell genes with weak overall predictability (cross-validated $R^2 < 0.45$) that were substantially more highly expressed in at least one disease group than in healthy donors (delta > 0.5 log2 units), the table reports the mean absolute Pearson correlation between methylation at the gene's top 50 highest-weight model CpGs and measured expression, computed separately in healthy donors and in the gene's own highest-expression adequately-powered diagnosis group (n >= 14: acute leukemia, healthy donors, other hematologic malignancies, lung cancer, and plasma cell disorder; chronic lymphocytic leukemia and lymphoma were underpowered and excluded). Columns: gene; healthy_coupling (mean absolute Pearson r in healthy donors); hiexpr_group (diagnosis group with highest mean expression among adequately-powered groups); hiexpr_mean_expr (mean log2 expression in that group); hiexpr_coupling (mean absolute Pearson r in that group); coupling_gain (hiexpr_coupling minus healthy_coupling); pattern (TRUE if coupling_gain > 0). Twelve of 15 genes show a positive coupling gain (binomial test against a null of 50%, p = 0.035).

**Supplementary Data S9. PBMC-retrained model performance compared to T cell models for a representative panel of 12 T cell genes.** Per-gene comparison of model performance between models trained on purified T cell methylation (T cell models) and models retrained directly on whole-PBMC methylation (PBMC-retrained models), evaluated on matched held-out test sets for a curated 12-gene representative panel spanning robust lineage markers (*CD8A*, *KLRG1*, *GZMB*, *TBX21*, *RUNX3*, *EOMES*), composition-sensitive markers (*CD3E*, *SPI1*), and additional comparison genes (*FOXP3*, *IL6*, *CD274*, *PDCD1*). Columns include: gene, T cell model cross-validated $R^2$, T cell model held-out test $R^2$, PBMC-retrained model held-out test $R^2$, number of CpGs selected by the PBMC model, number of stable CpGs in the T cell model, number of CpGs overlapping between the two models, percent overlap, and performance tier. The median CpG overlap between T cell and PBMC models across this panel is approximately 1.5%, confirming that the same lineage signal can be encoded by many alternative CpG combinations. This panel was selected to represent the range of transfer behaviors described in the main text and Fig. 5; results should not be generalized as performance estimates for all 77 genes.

**Supplementary Data S10. Composition-adjusted incidence associations for 76 (PBMC-weighted) or 77 (T cell-weighted) gene scores across 12 diseases in MGB/Harvard whole blood.** Full-adjusted Cox proportional hazards results (age, sex, and 11-cell EpiDISH composition adjustment, dropping neutrophils as the reference) for 76 evaluable gene scores in the PBMC-weighted model (CD80 excluded due to near-zero variance) and 77 gene scores in the T cell-weighted model (CD80 retained; its predicted score showed non-degenerate variance in this model) across 12 incident disease outcomes in the MGB/Harvard cohort (n = 4,250; median follow-up 8.6 years). Results are provided for both the T cell-weighted and PBMC-weighted model sets. Each row represents one gene-disease-model combination. Columns: gene, model (PBMC or

T cell), disease, n_at_risk, OR (odds ratio per standard deviation of predicted score), CI_low and CI_high (95% confidence interval bounds), p_value, BH_FDR (Benjamini-Hochberg false discovery rate, computed separately within each model (912 tests for the PBMC-weighted model; 924 tests for the T cell-weighted model)).

**Supplementary Data S11. Composition-adjusted all-cause mortality associations for 76 (PBMC-weighted) or 77 (T cell-weighted) gene scores in MGB/Harvard whole blood.** Full-adjusted Cox proportional hazards results (age, sex, and 11-cell EpiDISH composition adjustment) for 76 evaluable gene scores in the PBMC-weighted model and 77 gene scores in the T cell-weighted model, predicting all-cause mortality in the MGB/Harvard cohort (n = 4,239; 717 deaths; median follow-up 8.6 years). Results are provided for both the T cell-weighted and PBMC-weighted model sets. Columns: gene, model (PBMC or T cell), n, HR (hazard ratio per standard deviation of predicted score), CI_low and CI_high (95% confidence interval bounds), p_value, BH_FDR (Benjamini-Hochberg false discovery rate, computed separately within each model (76 tests for the PBMC-weighted model; 77 tests for the T cell-weighted model)).
