## Supplementary Figures and legends for "Regulatory mode and enhancer architecture determine limits of blood DNA methylation as a molecular proxy"

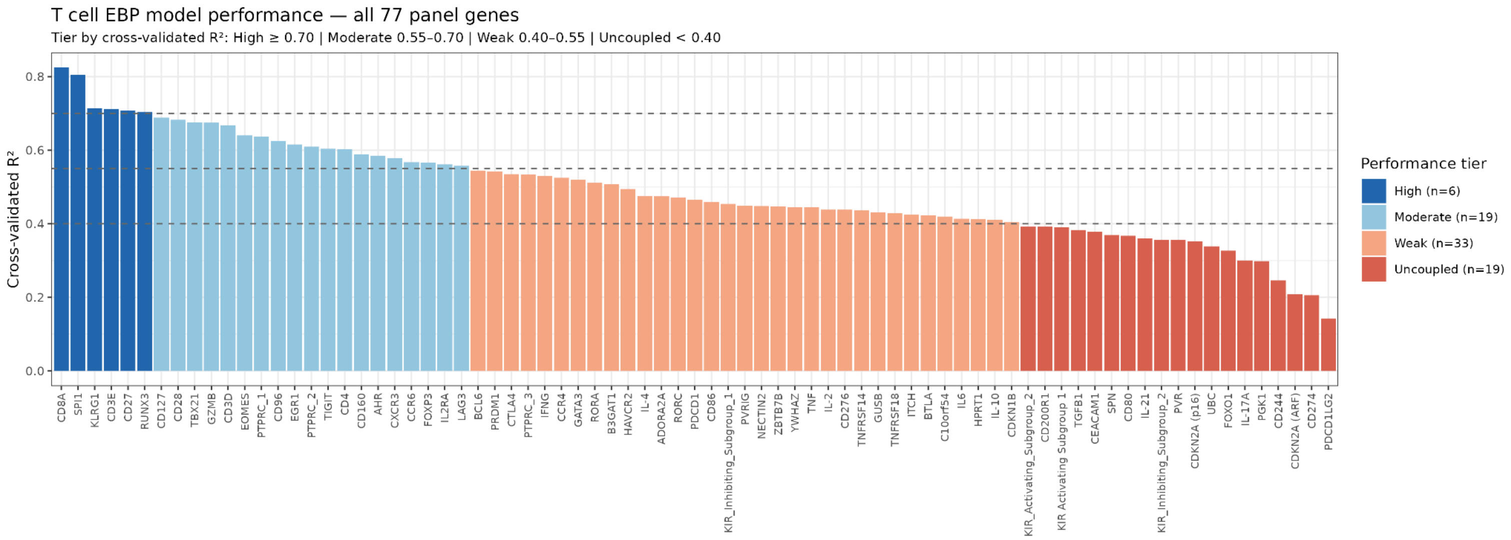


**Supplementary Figure S1. Cross-validated $R^2$ for all 77 T cell genes, ordered by performance.** Bar chart of cross-validated $R^2$ for each of the 77 T cell genes, ordered from highest to lowest. Bars are colored by performance tier (High ≥ 0.70, blue; Moderate 0.55 to 0.70, light blue; Weak 0.40 to 0.55, orange; Uncoupled < 0.40, red). The best model per gene was selected across four EWAS strategies (Pearson, Spearman, limma, duplicateCorrelation) by cross-validated $R^2$. $R^2$ values are from a post-alpha-selection, non-nested 10-fold cross-validation refit (alpha chosen once per gene via a separate 3-outer-fold/5-inner-fold search; Supplementary Note 2.17) on 333 matched blood samples. The two excluded genes (*GRAIL/RNF128* and *VTCN1/B7-H4*) are not shown; they were excluded because more than 25% of samples fell at the background-expression floor.

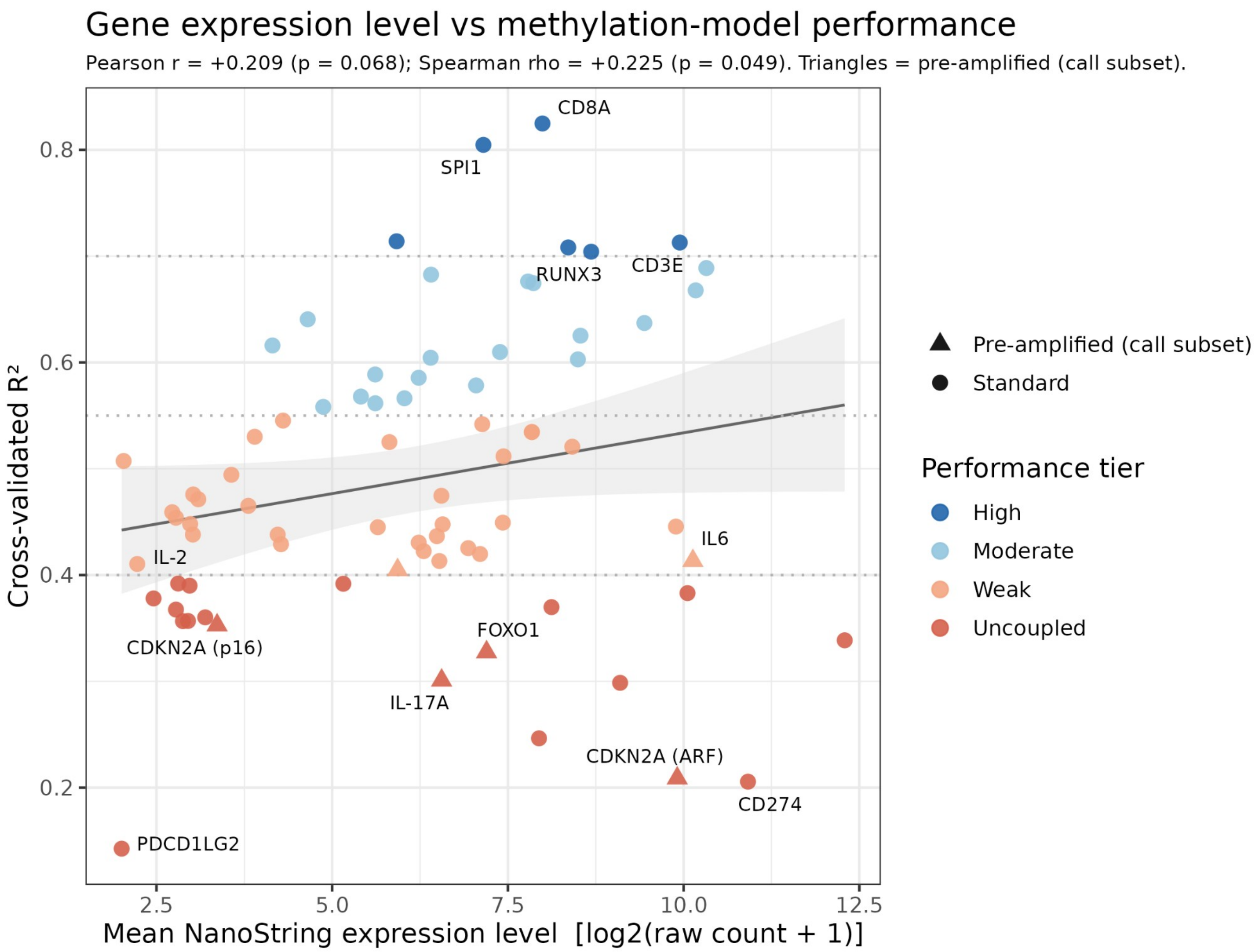


**Supplementary Figure S2. Mean NanoString expression is only weakly correlated with methylation-to-expression model performance.** Scatterplot of mean $\log_2$-normalized NanoString expression (x-axis) versus cross-validated $R^2$
(y-axis) for each of the 77 T cell genes. Each point is one gene; color indicates performance tier. Spearman rho = 0.23, p = 0.049, n = 77 genes. This weak association suggests that model performance is not primarily a consequence of expression abundance.

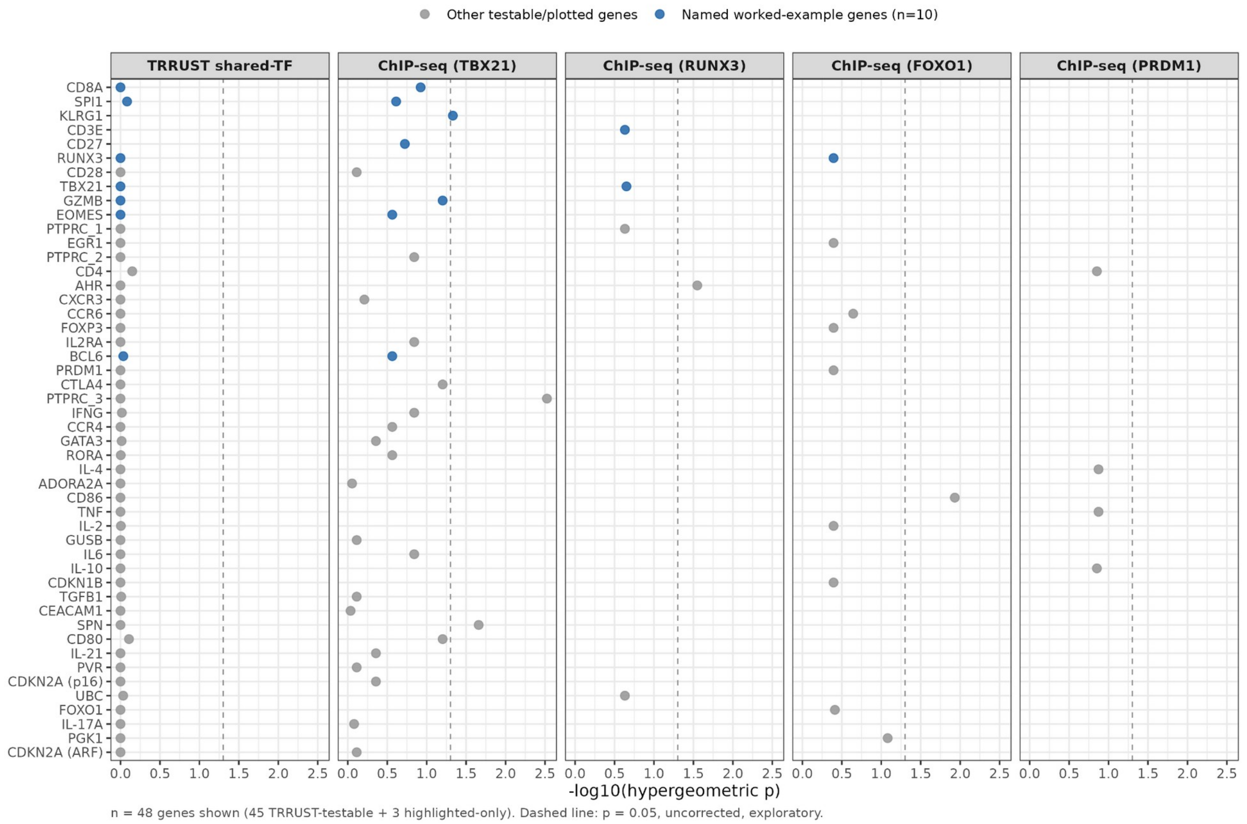


Supplementary Figure S3. Gene-by-gene transcription-factor enrichment across the TRRUST and ChIP-Atlas tests. Dot plot of -log10(hypergeometric p) for each gene's neighboring off-gene driver CpGs, faceted by test: TRRUST v2 shared-transcription-factor recurrence (leftmost panel) and ChIP-Atlas ChIP-seq peak overlap for four transcription factors (TBX21, RUNX3, FOXO1, PRDM1). Blue points indicate the 10 named worked-example genes discussed in the main text (CD8A, SPI1, KLRG1, CD3E, CD27, RUNX3, CD28, TBX21, GZMB, EOMES); gray points indicate the remaining testable or plotted genes (n = 48 genes shown: 45 TRRUST-testable plus 3 highlighted-only). The dashed vertical line marks p = 0.05 (uncorrected); none of the 45 TRRUST-testable genes reach this line in the leftmost panel, consistent with the absence of a shared canonical-regulator organization reported in the main text. The ChIP-seq panels show the small number of nominal hits (8 of 308 gene-by-transcription-factor tests) discussed in Methods and Supplementary Note 2 (Sections 2.15 and 2.16), four of which involve TBX21's unusually broad genome-wide binding background. Full per-gene values are provided in Supplementary Data S4.

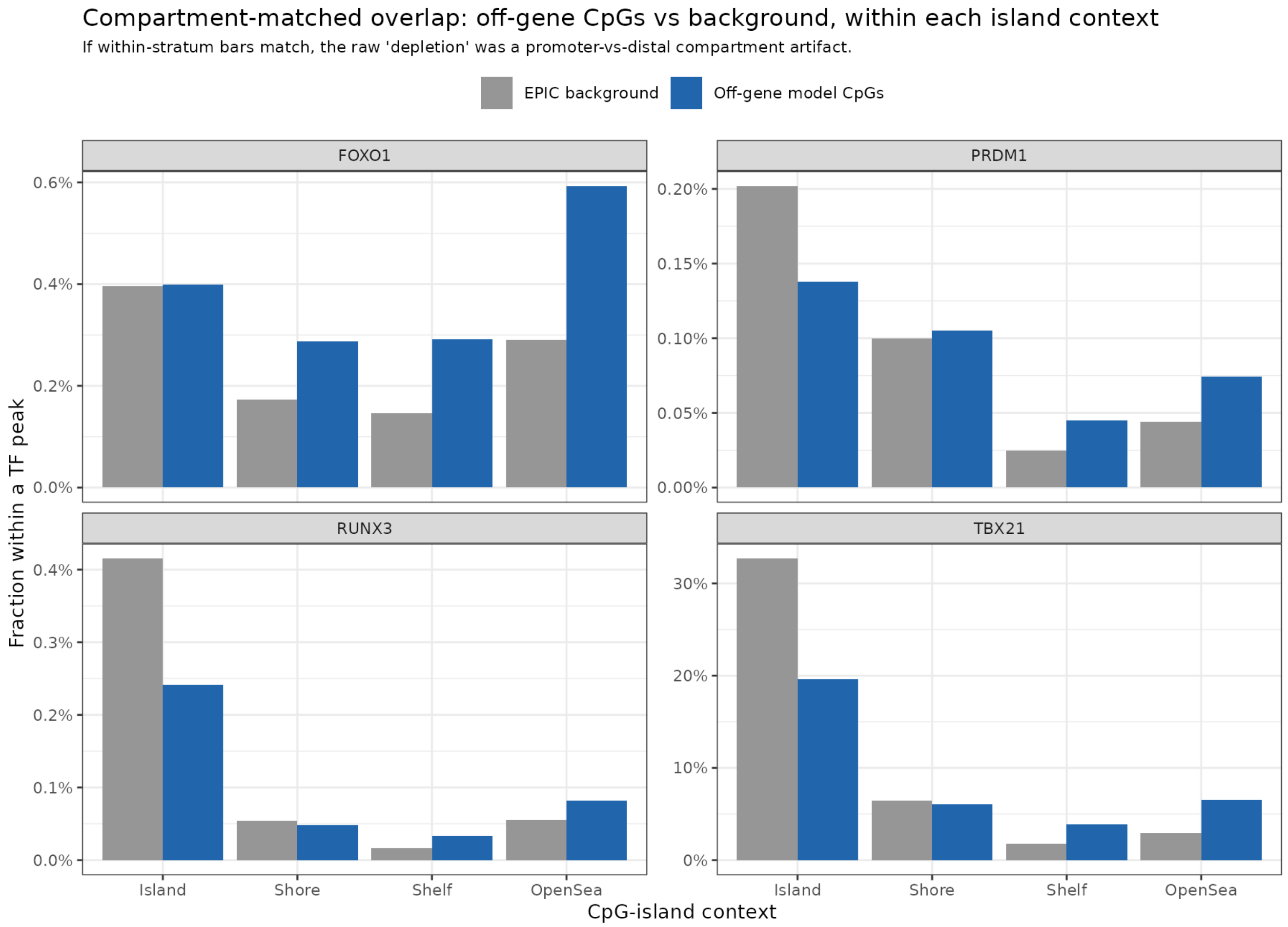


**Supplementary Figure S4. Compartment-matched overlap of off-gene model CpGs at transcription factor binding sites, within each CpG-island context.** For four transcription factors with available ChIP-seq data in immune cell types (*FOXO1, PRDM1, RUNX3,* and *TBX21*), bars show the fraction of CpGs overlapping a transcription factor peak for the full EPIC array background (grey) and the off-gene model CpGs (blue), stratified by CpG-island context (Island, Shore, Shelf, or OpenSea). Compartment matching addresses the confound that predictive CpGs are open-sea shifted while most transcription factor binding occurs at islands and promoters; if the raw depletion of off-gene CpGs at transcription factor peaks were a compartment artifact, within-stratum bars would match the background. For *TBX21* (primary Th1 T cell ChIP-seq), the within-island bar is lower for off-gene CpGs than background (approximately 20% versus 33%), confirming a genuine compartment-robust depletion of high-weight driver CpGs at the primary Th1 master regulator binding sites (Cochran-Mantel-Haenszel OR = 0.66, 95% CI 0.48 to 0.91, p = 0.01). *FOXO1, PRDM1,* and *RUNX3* show no consistent within-stratum depletion, consistent with their ChIP-seq being derived from non-primary T cell lines (GC-B cell, plasma cell, and NK-92, respectively) and therefore not informative for Th1 lineage CpG placement.

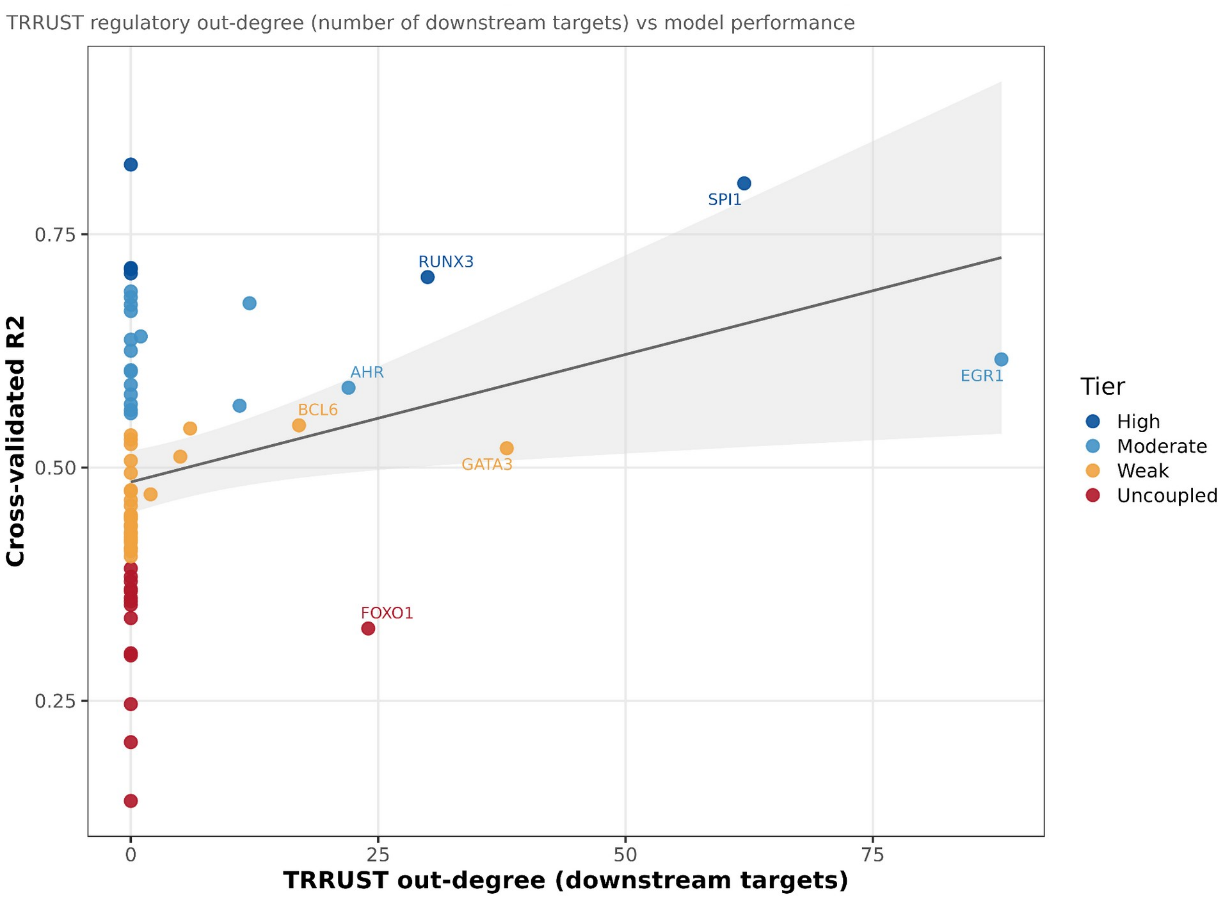


**Supplementary Figure S5. Transcription factor genes are better predicted from DNA methylation.** Scatterplot of TRRUST v2 regulatory out-degree (number of curated downstream targets, x-axis) versus cross-validated $R^2$ (y-axis) for each of the 77 T cell genes. Each point is one gene; color indicates performance tier; selected genes are labeled. Genes with higher regulatory out-degree, that is, genes that are themselves lineage master regulators, tend to be better predicted (Pearson r = 0.29, n = 77 genes). This is consistent with the lineage-versus-stimulus divide: transcription factors that define lineage programs are constitutively expressed in a lineage-dependent manner and are therefore more legible from DNA methylation. Their predictive CpGs are nonetheless not concentrated at canonical master-regulator binding sites (see Fig. 2e), indicating the effect reflects the regulatory mode of the gene rather than a direct transcription factor-binding mechanism.

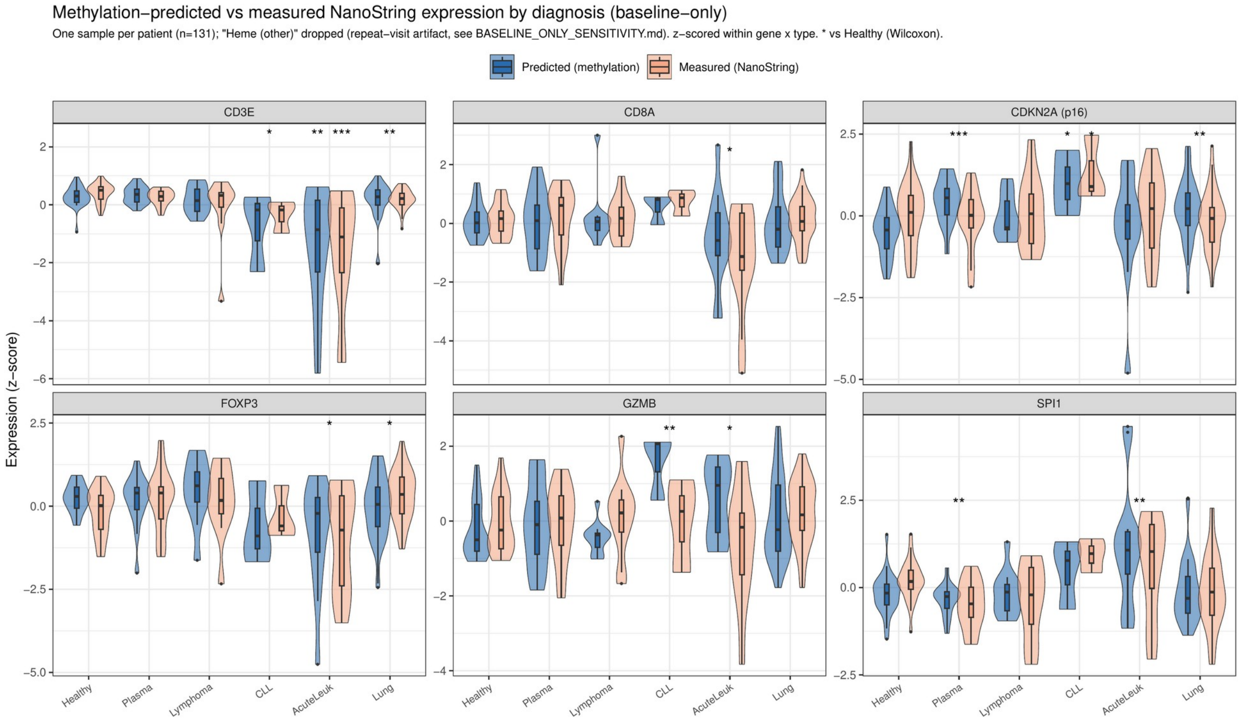


Supplementary Figure S6. Methylation-predicted versus directly measured NanoString expression by cancer diagnosis for representative genes, baseline-only. Violin plots comparing T cell-model methylation-predicted expression (blue) and directly measured NanoString expression (orange) for six representative genes (CD3E, CD8A, CDKN2A/p16, FOXP3, GZMB, and SPI1), restricted to one baseline (pre-treatment) sample per patient (n = 131) across the same five true diagnosis groups used in Fig. 4d and Supplementary Data S7 (plasma cell disorder, lymphoma, chronic lymphocytic leukemia, acute leukemia, and lung cancer); the pooled-sample “Heme (other)” category is excluded here for the same reason described in the main text (Methods) and Supplementary Note 2. Both scores are z-scored within the gene for comparable scales. Significance annotations indicate two-sided Wilcoxon rank-sum tests versus healthy donors (unadjusted; exploratory). The predicted scores track the measured expression across diagnosis groups for intrinsic lineage genes (CD3E, CD8A, GZMB, and FOXP3) while compressing within-group variance, consistent with the group-level concordance shown in Fig. 4a. SPI1 and p16 are included to illustrate the composition-driven (SPI1) and senescence-biology (p16) cases discussed in the main text. Models trained on the healthy Ohio State MAG cohort were applied without retraining to the baseline-only Burd/Rosko cancer subset (n = 131 of 228). n per diagnosis group as indicated.

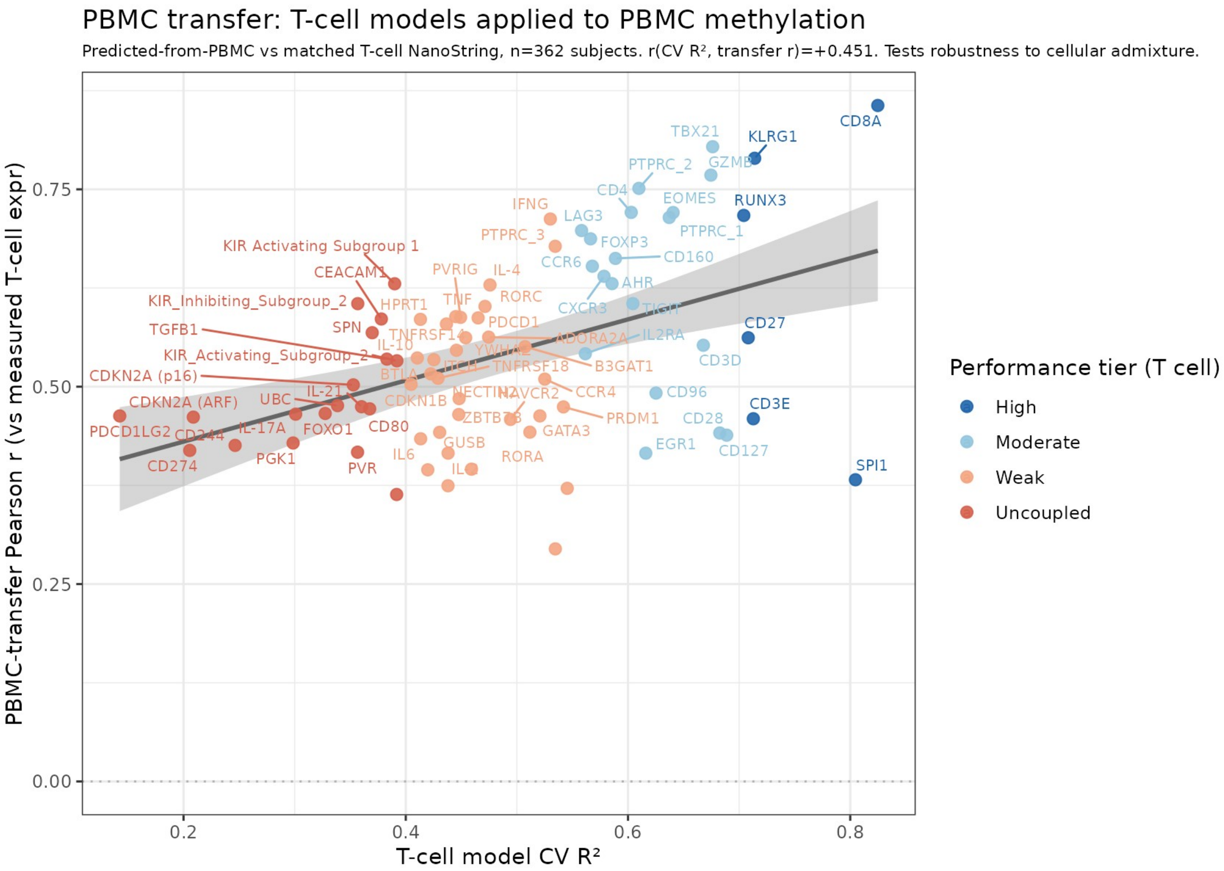


**Supplementary Figure S7. PBMC transfer Pearson correlation versus T cell model cross-validated $R^2$ for all 77 genes.** Scatterplot of PBMC transfer Pearson r (y-axis; correlation between methylation-predicted expression from PBMC methylation and directly measured purified-T-cell NanoString expression) versus T cell model cross-validated $R^2$ (x-axis) for each of the 77 T cell genes. Each point is one gene, labeled by name; color indicates performance tier. The positive correlation (r = 0.45, n = 77 genes) indicates that genes with stronger methylation-expression coupling in purified T cells also transfer more reliably to the PBMC admixture context. Intrinsic lineage genes (*CD8A, KLRG1, GZMB, TBX21,* and *EOMES*) cluster in the upper right; composition proxies (*SPI1*, *CD3E*) and stimulus-responsive genes cluster in the lower right. n = 362 matched PBMC-and-T-cell sample pairs.

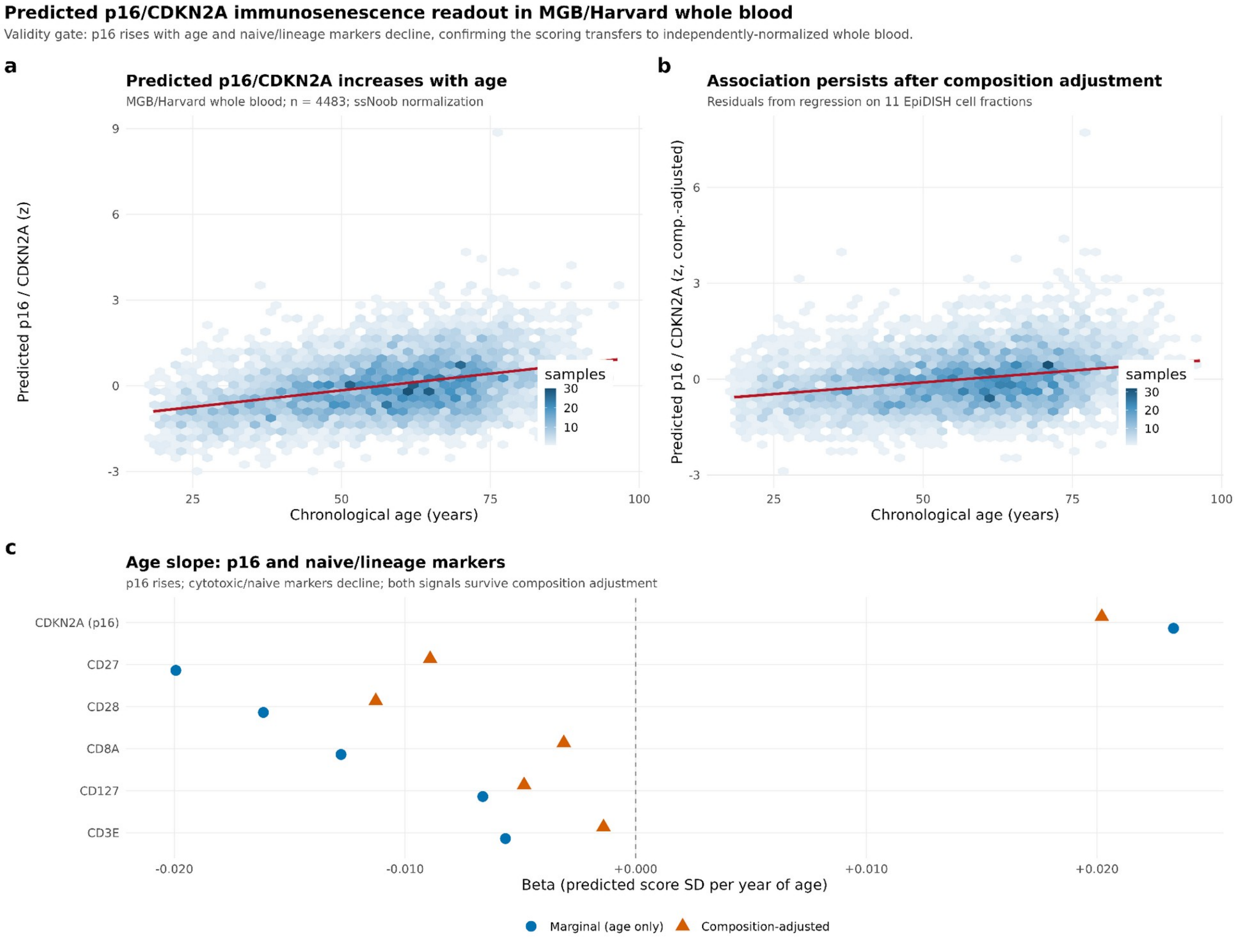


**Supplementary Figure S8. Predicted *p16/CDKN2A* immunosenescence readout in MGB/Harvard whole blood: validity gate. A)**, Predicted *p16/CDKN2A* score (standardized per SD) versus chronological age in the MGB/Harvard whole-blood cohort (n = 4,483; Illumina EPIC v1, ssNoob normalization). Each hexbin represents sample density; the red line is the least-squares regression (beta = +0.023 SD per year, p = 2.7 x $10^{-147}$, marginal). **B)**, Same association after partialing out 11 EpiDISH cell fractions (composition-adjusted residuals; beta = +0.020 SD per year, p = 5.0 x $10^{-105}$), confirming the age signal is not a cell-fraction artifact. **C)**, Age regression slope (beta per year) for *p16/CDKN2A* and five naive and lineage markers (*CD27, CD28, CD8A, CD127,* and *CD3E*), comparing marginal (blue circles) and composition-adjusted (orange triangles) estimates. *p16/CDKN2A* rises with age in both models; all lineage markers decline. The consistency of direction and magnitude across adjustment tiers confirms the scoring transfers to independently normalized whole blood and behaves biologically as expected.

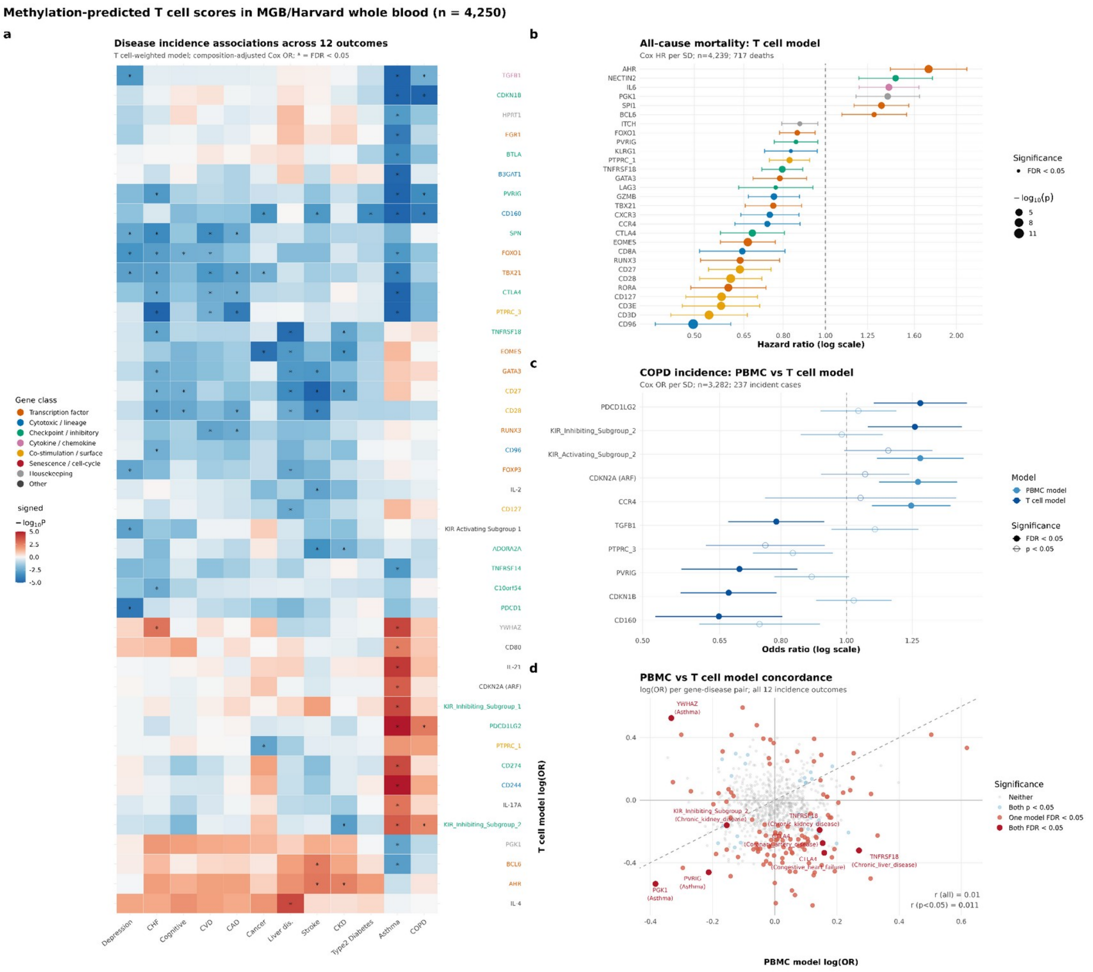


**Supplementary Figure S9. Methylation-predicted T cell scores in MGB/Harvard whole blood (n = 4,250). A)** Clustered heatmap of composition-adjusted incidence associations (Cox OR per standard deviation) across 77 gene scores and 12 diseases; T cell-weighted model, full adjustment (age, sex, 11-cell EpiDISH fractions). Color encodes signed -log10(P), capped at +/-5. Asterisks mark BH-FDR < 0.05 within the 924 tests. Gene labels are colored by functional class. Genes and diseases are ordered by hierarchical clustering on the T cell model association matrix. **B)** Forest plot of all-cause mortality associations from the T cell-weighted model (full composition adjustment; n = 4,239; 717 deaths; median follow-up 8.6 years). Each point is one gene score; filled circles indicate BH-FDR < 0.05, open circles p < 0.01. Point size encodes $-\log_{10}(P)$. Genes are ordered by hazard ratio. **C)** COPD incidence (n = 3,282 at risk; 237 incident cases) for both model sets. Each gene appears twice: light blue circle = PBMC model, dark blue triangle = T cell model. Filled symbols = BH-FDR < 0.05 within the full incidence scan; open symbols = p < 0.05. Genes shown if FDR < 0.05 in at least one model or p < 0.01 in both. **D)** Concordance scatter of PBMC model versus T cell model log(OR) for all 912 gene-disease incidence pairs. Color encodes significance status. The low overall correlation (r = 0.01) reflects complementary rather than redundant model readouts; directional agreement is higher among the most significant individual pairs (labeled).

#### SUPPLEMENTARY METHODS

To demonstrate deployment at population scale in the most accessible sample type, both the T cell-weighted and PBMC-weighted model sets were scored on the MGB/Harvard Biobank whole-blood cohort (Illumina EPIC v1; ssNoob normalization; n = 4,250 with complete phenotype and outcome data; 4,483 with age for the validity gate). Scoring used a memory-safe per-model M-value conversion implemented in the deployment bundle (score_beta_matrix.R); each predicted gene score was standardized to mean 0 and unit standard deviation within the analysis set (per-SD scaling), so all reported hazard ratios and odds ratios are per-SD effects. One gene score, CD80, showed near-zero variance in the PBMC-weighted model and was excluded from that model only, leaving 76 evaluable PBMC scores; CD80 was retained in the T cell-weighted model, where its predicted score showed non-degenerate variance, leaving all 77 T cell scores evaluable.

Composition was estimated by EpiDISH RPC with the cent12CT reference as above; all disease models below include 11 of the 12 cell fractions as covariates (neutrophils were dropped as the sum-to-one reference to avoid singularity). Mandatory composition adjustment was applied to every result reported from this cohort, because whole blood is granulocyte-dominated (approximately 60% neutrophils) and T cells comprise only approximately 15 to 25% of white cells; without adjustment, any association with immune or age-related disease can be wholly explained by cell-fraction differences across individuals.

As a validity gate applied before any disease analysis, predicted *p16/CDKN2A* was regressed on chronological age both marginally and after composition adjustment (linear regression, age as continuous predictor), and the naive and *CD8* lineage scores (*CD27, CD28, CD8A, CD127,* and *CD3E*) were checked for the expected age-related decline. This gate was required to pass before any disease analysis was interpreted.

Disease outcome data were obtained from structured EHR records linked to the MGB Biobank. Incident outcomes were defined as a new ICD-coded diagnosis occurring after the methylation collection date, excluding participants with a prevalent diagnosis at baseline. All-cause mortality was ascertained from hospital records and death registration. Twelve incident disease outcomes were tested (Asthma, COPD, Type 2 Diabetes, Coronary Artery Disease, CVD excluding stroke, Stroke, Congestive Heart Failure, Chronic Kidney Disease, Chronic Liver Disease, Cancer, Depression, and Cognitive Deficit), plus all-cause mortality. These outcomes were selected by the MGB collaborators (M.M., J.L.S.) based on availability and event count in the biobank; no pre-specified primary outcome was designated for the MGB analysis, which is treated as a discovery-and-illustration deployment rather than a confirmatory study.

For all-cause mortality and each incident disease, composition-adjusted Cox proportional-hazards models were fit using the survival package v3.7.0 (Therneau 2024 [CRAN survival]) with time-to-event as the follow-up from collection date to event or censoring. Cox PH was chosen because it appropriately handles right-censored follow-up, is the standard model for incident disease in biobank EHR cohorts, and yields hazard ratios (HRs) directly interpretable as per-SD effects on the instantaneous event rate. The proportional-hazards assumption was not formally tested at this scale (912 PBMC models and 924 T cell models); violations would attenuate effect estimates toward the null. For each outcome, models were fitted three ways: unadjusted (score only), semi-adjusted (score plus age and sex), and fully adjusted (score plus age, sex, and 11 EpiDISH cell fractions); only fully adjusted results are reported in the main text. Cross-sectional (prevalent) associations used logistic regression with the same covariate structure.

Benjamini-Hochberg false discovery rate (FDR) correction was applied separately within each model set, with a significance threshold of FDR less than 0.05: 912 gene-disease tests (76 genes times 12 diseases) for the PBMC-weighted model, and 924 gene-disease tests (77 genes times 12 diseases) for the T cell-weighted model. Separate FDR correction for mortality was applied independently (76 tests for the PBMC-weighted model, 77 tests for the T cell-weighted model). No correction was applied across the PBMC and T cell model sets; the two models were treated as parallel discovery analyses rather than replication.

To characterize the concordance and complementarity of the two model sets, Pearson correlation of log(OR) was computed across all 912 gene-disease incidence pairs for the 76 genes common to both model sets (CD80, evaluable only in the T cell-weighted model, was not included in this comparison). Low overall concordance (r = 0.01) indicates complementary rather than redundant readouts, consistent with the 1.5% CpG overlap between T cell and PBMC-retrained models established in the admixture analysis.

**Supplementary Note 1: Software, Package Installation, and External Data Sources**

Genome build: hg19 / GRCh37 throughout. Any GRCh38 source was lifted to hg19 via the UCSC hg38ToHg19.over.chain before use.

### Part 1. R environment and package installation

All analyses were performed in R version 4.4.0 (R Core Team 2024). The following commands reproduce the package environment used in the study. Bioconductor packages require BiocManager; CRAN packages use install.packages().

```
# Bioconductor (version 3.20)
if (!requireNamespace("BiocManager", quietly = TRUE))
    install.packages("BiocManager")
BiocManager::install(version = "3.20")
BiocManager::install(c(
  "minfi",                                        # v1.52.1  -- array preprocessing
  "IlluminaHumanMethylationEPICanno.ilm10b4.hg19",   # v0.6.0   -- EPIC manifest
  "limma",                                        # v3.62.1  -- linear models,
duplicateCorrelation
  "missMethyl",                                   # v1.40.3  -- CpG-bias-aware enrichment
(gometh)
  "clusterProfiler",                              # v4.14.6  -- GO/KEGG enrichment
  "org.Hs.eg.db",                                 # v3.20.0  -- human gene annotation
  "TxDb.Hsapiens.UCSC.hg19.knownGene",            # v3.2.2   -- nearest-annotated-gene
mapping
                                                     #          (500 kb window, TF-module
test)
  "ReactomePA",                                   # v1.50.0  -- Reactome enrichment
  "reactome.db",                                  # v1.89.0  -- Reactome database
  "GenomicRanges",                                # v1.58.0  -- genomic interval
operations
  "rtracklayer",                                  # v1.66.0  -- BED/bigWig import,
liftOver
  "EpiDISH",                                      # v2.22.0  -- reference-based cell
deconvolution
  "DESeq2",                                       # v1.46.0  -- expression normalization
  "RUVSeq"                                        # v1.40.0  -- remove unwanted variation
))

# CRAN
install.packages(c(
  "glmnet",      # v5.0       -- elastic net models
  "survival",    # v3.7.0   -- Cox proportional hazards
  "msigdbr",     # v25.1.1  -- MSigDB gene sets (Hallmark, C7 immunologic)
  "data.table",  # v1.16.4  -- fast data manipulation
  "ggplot2",     # v3.5.1   -- visualization
  "patchwork",   # v1.3.0   -- multi-panel figure assembly
  "ggrepel",     # v0.9.6   -- non-overlapping labels
  "ragg",        # v1.3.3   -- high-resolution PNG/TIFF output
  "scales",      # v1.3.0   -- axis formatting
  "dplyr",       # v1.1.4   -- data manipulation
  "tidyr"        # v1.3.1   -- data reshaping
```

```
))
```

##### Key package citations

| Package | Version | Reference |
|---|---|---|
| minfi | 1.52.1 | Aryee et al. 2014, PMID 24478339 |
| limma | 3.62.1 | Ritchie et al. 2015, PMID 25605792 |
| glmnet | 5.0 | Friedman et al. 2010, PMID 20808728 |
| EpiDISH | 2.22.0 | Teschendorff et al. 2017, PMID 28193155 |
| clusterProfiler | 4.14.6 | Wu et al. 2021, PMID 34557778 |
| missMethyl | 1.40.3 | Phipson et al. 2016, PMID 26424855 |
| ReactomePA | 1.50.0 | Yu and He 2016, PMID 26661513 |
| GenomicRanges | 1.58.0 | Lawrence et al. 2013, PMID 23950696 |
| DESeq2 | 1.46.0 | Love et al. 2014, PMID 25516281 |
| RUVSeq | 1.40.0 | Risso et al. 2014, PMID 25150836 |
| survival | 3.7.0 | Therneau and Grambsch 2000 |
| msigdbr | 25.1.1 | Liberzon et al. 2015 |
| TxDb.Hsapiens.UCSC.hg19.knownGene | 3.2.2 | Bioconductor annotation package, no primary citation |

**Second package-version snapshot, disclosed separately (2026-07-04 age-confound reanalysis and Figure 4 reconstruction).** This analysis ran in a later package environment than the primary pipeline above for two plotting/data packages; both snapshots are reported rather than merged into one falsely-uniform list, because we confirmed the versions genuinely differ (verified directly against the execution environment on 2026-07-04): R 4.4.0; dplyr 1.2.1 (primary pipeline: 1.1.4); tidyr 1.3.2 (primary pipeline: 1.3.1); ggplot2 4.0.3 (primary pipeline: 3.5.1); patchwork 1.3.2 (primary pipeline: 1.3.0); magick 2.8.5 (new to this analysis, used to assemble the corrected Figure 4); EpiDISH 2.22.0 (unchanged). This reanalysis did not refit any statistical model with a different package version – the linear models in Part 2 below used the same R and EpiDISH versions as the primary pipeline – the version difference affects only figure rendering.

### Part 2. External datasets and reference resources

#### 2.1 Cohort methylation data (internal – not publicly available)

| Dataset | Description | Normalization | N |
|---|---|---|---|
| Training (T cell) | Ohio State MAG cohort, purified T cells, Illumina EPIC v1 | Reference-anchored functional normalization (global PCA from ~22,000 EPIC v1 samples) followed by RCP probe-type bias correction and detection p-value masking | 333 samples |
| Cancer validation | Burd/Rosko cohort (OSU), Illumina EPIC v1 | Functional normalization (preprocessFunnorm) followed by RCP (betaRCP) | 228 samples |
| PBMC admixture | Ohio State MAG cohort, whole PBMC, Illumina EPIC v1 | Same reference-anchored functional normalization plus RCP as training cohort | 385 samples |
| MGB whole blood | Mass General Brigham Biobank, Illumina EPIC v1 | ssNoob (Triche et al. 2013) | 4,250 analyzed |

Cohort methylation data are not publicly deposited due to participant privacy; model weights and deployment scripts are available from the corresponding author after a signed Data Use Agreement (not simply "upon reasonable request" – the training data cannot be shared without one).

#### 2.2 Array annotation

| Resource | Version | Source | Build | Use |
|---|---|---|---|---|
| Illumina EPIC manifest | IlluminaHumanMethylationEPICanno.ilm10b4.hg19 v0.6.0 | Bioconductor | hg19 | CpG-to-gene, region, and CpG-island context annotation throughout |

#### 2.3 Chromatin state maps – Roadmap Epigenomics

Source: https://egg2.wustl.edu/roadmap/data/byFileType/chromhmmSegmentations/ChmmModels/coreMarks/jointModel/final/

Citation: Roadmap Epigenomics Consortium, Nature 2015, PMID 25693563

Model: 15-state ChromHMM coreMarks mnemonics, hg19

Download date: May 2026 (resting T cell states); June 2026 (activated states)

| Epigenome ID | Cell type | Condition | File |
|---|---|---|---|
| E043 | CD4+ naive T (primary) | Resting | E043_15_coreMarks_mnemonics.bed.gz |
| E044 | CD4+ memory T (primary) | Resting | E044_15_coreMarks_mnemonics.bed.gz |
| E047 | CD8+ naive T (primary) | Resting | E047_15_coreMarks_mnemonics.bed.gz |
| E041 | Primary Th cells | PMA-I stimulated | E041_15_coreMarks_mnemonics.bed.gz |
| E042 | Primary Th17 cells | PMA-I stimulated | E042_15_coreMarks_mnemonics.bed.gz |

#### 2.4 Canonical TF ChIP-seq – ENCODE

Source: https://www.encodeproject.org

Build: GRCh38 peaks lifted to hg19 with UCSC liftOver (hg38ToHg19.over.chain)

Download date: May 2026

| TF | Experiment | File accession | Biosample | Cell type |
|---|---|---|---|---|
| CTCF | ENCSR470KCE | ENCFF858TLX | CD4+ naive T | Primary |
| CTCF | ENCSR116AKQ | ENCFF758CQW | CD8+ naive T | Primary |
| ETS1 | ENCSR000BKA | ENCFF332PGQ | GM12878 | Lymphoblastoid line |
| SP1 | ENCSR000BKQ | ENCFF076YZO | K562 | CML line |

#### 2.5 Lineage master regulator TF ChIP-seq – ChIP-Atlas

Source: https://chip-atlas.dbcls.jp

Stringency: q less than $1 \times 10^{-5}$ (bed05 threshold)

Citation: Oki et al. 2018, EMBO Rep, PMID 30413482

Build: hg19 (pre-called)

Download date: June 2026

| TF | Cell type | Primary vs line | SRX accessions |
|---|---|---|---|
| TBX21 | Th1 (primary CD4, in vitro differentiated) | Primary T cell | SRX092313, SRX1799591, SRX1799593, SRX735289 |
| RUNX3 | NK-92 | Cell line (NK lymphoma) | SRX19719536, SRX19719537, SRX19719540, SRX19719541 |
| PRDM1 | U-266 | Cell line (multiple myeloma) | SRX3070540 |
| FOXO1 | Germinal-center B (tonsil) | Primary B cell | SRX1012536, SRX1012537 |
| EOMES | None available | No immune ChIP-seq in ChIP-Atlas, ENCODE, or ReMap | -- |

Note: Only TBX21 has primary T cell ChIP-seq data. RUNX3, PRDM1, and FOXO1 peaks are from non-T-cell lines and are used for illustration only. EOMES is absent entirely. This same dataset is reused as a second, independent evidence source for the gene-by-gene transcription-factor-module test (section 2.10, below).

#### 2.6 Regulatory network – TRRUST v2

Source: https://www.grnpedia.org/trrust/

File: trrust_rawdata.human.tsv

Citation: Han et al. 2018, Nucleic Acids Research, PMID 29087512

Download date: May 2026

Content: Manually curated human transcription factor to target-gene regulatory relationships (direction: activation/repression; evidence: published literature)

#### 2.7 Pathway and ontology databases

| Collection | Package | Version | Notes |
|---|---|---|---|
| Gene Ontology (BP, MF, CC) | org.Hs.eg.db via clusterProfiler | v3.20.0 | All three ontologies tested |
| KEGG | clusterProfiler::enrichKEGG | v4.14.6 | Organism hsa |
| Reactome | ReactomePA with reactome.db | v1.50.0 / v1.89.0 | Yu and He 2016 |
| MSigDB Hallmark (H) | msigdbr | v25.1.1 | 50 hallmark gene sets |
| MSigDB C7 immunologic | msigdbr | v25.1.1 | Immune perturbation signatures |

#### 2.8 Cell-type deconvolution reference

| Resource | Source | Use |
|---|---|---|
| EpiDISH cent12CT reference matrix | Built into EpiDISH v2.22.0 (Luo et al. 2023, PMID 37525279) | 12-cell blood composition estimation (CD4 naive/memory, CD8 naive/memory, Treg, B naive/memory, NK, Mono, Neu, Eos, Baso) using robust partial correlation (RPC method) |

Note: distinct from Salas et al. 2022, PMID 35140201 (Nat Commun 13:761, "Enhanced cell deconvolution of peripheral blood using DNA methylation for high-resolution immune profiling"), the IDOL 12-cell deconvolution library paper cited elsewhere in this manuscript as a separate precedent reference. Both are correctly cited; they are not interchangeable.

#### 2.9 UCSC liftOver chain

Source: https://hgdownload.soe.ucsc.edu/goldenPath/hg38/liftOver/hg38ToHg19.over.chain.gz
Use: Lifting ENCODE GRCh38 ChIP-seq peaks to hg19

#### 2.10 Nearest-gene mapping for the gene-by-gene transcription-factor-module test

Resource: TxDb.Hsapiens.UCSC.hg19.knownGene and org.Hs.eg.db (Bioconductor annotation packages, versions as in Part 1).

Use: for each gene's top 20 highest-absolute-weight off-gene driver CpGs, mapping to the nearest annotated gene within a 500 kb window (hg19), the first genomic-distance nearest-gene lookup used in this study (all other CpG-to-gene annotation in this manuscript uses only the EPIC manifest's gene-body/promoter overlap annotation, which is silent for CpGs with no direct gene-body overlap). Promoter overlap for the ChIP-seq sensitivity check (section 2.5) used a transcription-start-site window of plus or minus 2 kb. Download/build date: 2026-07-03, alongside the gene-by-gene TF-module analysis itself.

**Supplementary Note 2: Extended Methods for Cancer-Cohort Validation and Off-Gene Regulatory Robustness Tests**

#### 2.11 Cell-composition reference panel and CD3-enrichment calibration (cancer cohort)

The EpiDISH cent12CT reference matrix used throughout this manuscript provides reference profiles for 12 blood cell types: four CD4 T cell subsets (naive, memory), two CD8 T cell subsets (naive, memory), two B cell subsets (naive, memory), NK cells, monocytes, eosinophils, and basophils. Sorted-T training samples were deconvolved under this same reference to establish a calibration baseline: the resulting T cell fraction of approximately 83% (versus 100% expected for sorted T cells) was consistent across samples and reflects a systematic reference under-call rather than true contamination.

This 83% calibration baseline was compared, diagnosis by diagnosis, against the Burd/Rosko cancer cohort's own deconvolved composition. Four of six disease groups matched the baseline closely (T cell fraction 82.5% to 84.8%, B cell fraction 2.7% to 4.2%), consistent with the intended CD3 enrichment. Two groups deviated: chronic lymphocytic leukemia (T cell 70.9%, B cell 16.0%, n = 5) and acute leukemia (T cell 70.1%, B cell 5.9%, n = 22), consistent with residual circulating malignant cells surviving the enrichment step in diseases where the circulating leukocyte pool is itself dominated by the malignant clone. This diagnosis-specific residual composition, not a general whole-blood composite, is what the disease-association composition adjustment (main text Methods) controls for.

#### 2.12 Composition-covariate rank deficiency (eosinophils and basophils)

In the cancer-cohort disease-association models, CD8_total and CD4_total are the sums of the naive and memory subsets of each lineage from the 12-cell cent12CT deconvolution, and Treg, B_total (Bnv + Bmem), NK, Mono, and Neu are as returned by EpiDISH. Eosinophils and basophils were excluded from the composition covariates because their fractions are near zero in most samples and their inclusion produced rank-deficient models (near-singular design matrix). This matches the covariate set used identically in the PBMC admixture-transfer composition control (main text Methods). The composition adjustment used throughout this cohort's disease-association testing is therefore 7 grouped cell-fraction covariates, not the full 12 raw cent12CT columns.

#### 2.13 Multiple-testing rationale for the cancer-cohort disease-association family

The 40-test cancer-cohort disease-association family (8 genes by 5 diagnosis groups, evaluated under three nested models: unadjusted, age-adjusted, and fully adjusted) was not corrected for multiple testing, nor was the original p16-only Kruskal-Wallis and Wilcoxon comparison. This choice is deliberate and is stated explicitly here rather than left implicit: the cancer cohort's smallest diagnosis groups (chronic lymphocytic leukemia, n = 5; lymphoma, n = 14) make this an underpowered, hypothesis-generating screen rather than a confirmatory test, and applying FDR correction to a test family of this size and power would not meaningfully change which associations are trustworthy. Instead, the full three-model comparison for every test is reported directly (Supplementary Data S7), so that readers can judge robustness from the age- and composition-adjustment pattern rather than from a corrected p-value alone.

#### 2.14 Detectability and cell-fraction mechanism test (coupling shift)

As a complementary test of whether genes with weak overall predictability are truly uncoupled from methylation, or are instead limited by low expression variance and cell-fraction dilution in the training cohort, we identified 15 candidate genes with weak cross-validated predictability ($R^2$ less than 0.45) that were nonetheless substantially more highly expressed in at least one cancer-cohort diagnosis group than in healthy donors (delta greater than 0.5 $\log_2$ units in measured NanoString expression). For each candidate gene, we computed the mean absolute Pearson correlation between M-values at the gene's top 50 highest-absolute-weight model CpGs and measured expression, separately within each diagnosis group with at least 14 samples (acute leukemia, healthy, other hematologic malignancies, lung cancer, and plasma cell disorder; chronic lymphocytic leukemia, n = 5, and lymphoma, n = 14, were treated as inadequately powered and excluded from this specific test, as the boundary $n \geq 14$ criterion is the least stable given the additional 50-CpG-by-sample requirement).

We then compared each gene's coupling strength in healthy donors to its coupling strength in its own highest-expression, adequately powered group. Coupling strengthened in the higher-expression group for 12 of 15 genes (two-sided exact binomial test against a null of 50%, p =

0.035); the three exceptions were CEACAM1, BTLA (for which no disease group exceeded healthy-donor expression, so no comparison was possible), and PDCD1LG2. This is a single, pre-specified binomial test and was not part of a multiple-testing family requiring correction. Full per-gene results, including the selection criterion, the healthy and highest-expression-group coupling values, and the gain for each gene, are provided in Supplementary Data S8.

#### 2.15 Gene-by-gene shared-transcription-factor test (TRRUST v2)

Because the pooled recurrence test (main text Methods) treats all 77 genes' driver CpGs as one interchangeable set, we additionally tested, gene by gene, whether each gene's own extragenic driver CpGs are organized around that gene's own known regulators, rather than asking only whether driver CpGs recur across different genes' models. For each gene, the top 20 highest-absolute-weight off-gene driver CpGs were mapped to their nearest annotated gene within 500 kb (hg19, TxDb.Hsapiens.UCSC.hg19.knownGene and org.Hs.eg.db); genes with no off-gene driver CpG within this window were excluded. The gene's own curated upstream transcription-factor regulators were looked up in TRRUST v2, and each neighboring gene's curated upstream regulators were looked up the same way.

A gene was classified as testable only if it had at least one curated TRRUST regulator and at least one neighboring gene within the 500-kb window; 32 of 77 genes had no curated TRRUST regulator and were excluded as untestable, leaving 45 testable genes. For each testable gene, the number of neighboring genes sharing at least one transcription factor with the index gene was compared to chance by a one-sided hypergeometric test, with the population defined as all genes annotated as a Target anywhere in TRRUST v2, the number of population genes sharing a factor with the index gene as the number of successes, and the count of the index gene's own neighboring genes as the sample size. None of the 45 testable genes reached nominal significance (smallest $p = 0.71$), including all 7 testable genes among a set of 10 named lineage and regulatory genes examined in detail during manuscript review (CD8A, RUNX3, TBX21, GZMB, EOMES, SPI1, BCL6; the remaining 3 of the 10 — KLRG1, CD3E, and CD27 — had no curated TRRUST regulator and were untestable).

#### 2.16 Independent ChIP-Atlas evidence and genome-wide sensitivity check

Because TRRUST v2 curation is sparse for T cell surface and lineage genes (for example, it contains no RUNX3-to-CD8A edge, even though RUNX3 is an established CD8-lineage regulator in the primary literature; CD8A's only curated TRRUST regulators are ETS1 and GATA3), we added a second, independent evidence source for the specific case examined in detail (CD8A driver-CpG neighbors and RUNX3): ChIP-Atlas ChIP-seq peak overlap (chip-atlas.dbcls.jp; the same TBX21, RUNX3, PRDM1, and FOXO1 peak sets used in the Transcription-factor binding-site overlap section, main text Methods) at each neighboring gene's promoter (transcription start site plus or minus 2 kb). None of CD8A's 18 driver-CpG neighboring gene promoters were RUNX3-bound (0 of 18), against a genome-wide background RUNX3 promoter-binding rate of 1.3% (one-sided hypergeometric p, population and background defined analogously to the TRRUST test above but using genome-wide promoter counts and binding counts in place of TRRUST-curated relationships).

Extending this ChIP-seq sensitivity check across all 77 genes and all 4 available transcription factors (308 gene-by-transcription-factor tests), 8 of 308 reached nominal p less than 0.05; 4 of these 8 involved TBX21, whose ChIP-Atlas pooled peak set covers 55.7% of all gene promoters genome-wide, an unusually broad background that likely reflects lenient pre-called peak calling (q less than $1 \times 10^{-5}$ across pooled experiments) rather than genuine TBX21 specificity, and these 4 hits are interpreted with that caveat; the remaining 4 (RUNX3- and FOXO1-linked, background promoter-binding rates 0.75% to 2.5%) rely on non-primary-T-cell ChIP-seq (NK-92, B-cell, and myeloma lines, as noted in the Transcription-factor binding-site overlap section) and are reported as suggestive rather than confirmatory. This gene-by-gene test and the pooled recurrence test (main text Methods) ask related but distinct questions — whether specific known regulators organize each gene's off-gene signal, versus whether the same CpGs recur across different genes' models — and reach the same conclusion by different methods: neither finds evidence that off-gene driver CpGs are organized around a small set of shared or canonical regulators. Full per-gene results, including the TRRUST testability and hypergeometric-p columns and the ChIP-seq sensitivity columns described above, are provided in Supplementary Data S4.

#### 2.17 Elastic-net hyperparameters and cross-validation procedure

Predictive models were elastic nets fit with glmnet v5.0 [31]. The elastic net was chosen because it combines L1 (lasso) and L2 (ridge) penalties, enabling sparse feature selection while handling the collinearity inherent in genome-wide methylation data. The outer cross-validation used three folds defined at the subject level (seed 42, assigned via a stratified split so that all visits from the same subject appeared in the same fold), ensuring that generalization estimates reflect subject-level rather than visit-level held-out performance. The alpha mixing parameter was tuned over a grid of six values (0.01, 0.1, 0.25, 0.5, 0.75, 1.0) – ranging from near-ridge to lasso – with mean cross-validated MSE averaged across the three outer folds and a single best-performing alpha chosen from that pooled mean, rather than an independent alpha winner selected per fold. For this alpha-tuning step and for calling stable CpGs, the top-50,000-CpG feature set was recomputed fresh within each outer fold’s training portion only, so neither the selected alpha nor the stable-CpG calls are affected by feature-selection leakage. The cross-validated $R^2$ quoted in the main text, however, is computed differently: the top-50,000-CpG feature set was instead selected once from the full training data, and the model was then refit at the best alpha with a 10-fold inner cross-validation for lambda selection only (lambda.min criterion) on that fixed feature set. Because the feature-selection step for this number is not itself repeated within each of its own 10 folds, it carries feature-selection leakage relative to those folds and should be read as a training-cohort performance estimate rather than a fully leakage-free one; the held-out test $R^2$ computed on the 20% of samples never used in any selection or fitting step is not subject to this leakage and is reported separately for the on-gene-restricted retraining comparison (Section 2.18) and the PBMC cross-biospecimen retraining comparison (Section 2.22). Cross-validated $R^2$ was defined as max(0, 1 minus the ratio of cross-validated MSE to the variance of the outcome), with the floor at zero preventing negative values from inflating the apparent performance of uninformative genes.

#### 2.18 On-gene versus off-gene CpG classification and on-gene-restricted retraining

To determine where each model's predictive weight falls relative to the target gene, every stable CpG in each model was classified as on-gene (annotated to the target gene's own promoter or body in the hg19 EPIC manifest) or off-gene (all other locations: other genes' regulatory regions, or intergenic and distal sites), and the weight share of each class was computed as the sum of absolute elastic-net coefficients for that class divided by the total. To test whether off-gene CpGs carry information that on-gene CpGs lack – rather than simply being more numerous – we retrained, for each gene, an elastic net restricted to on-gene CpGs only and compared its held-out test $R^2$ to the genome-wide top-50,000 model on the identical subject-level 80/20 split. This comparison was performed at a fixed alpha of 0.5 and a single seed; it is an illustrative control, not an optimized model.

#### 2.19 Functional enrichment: database parameters and package versions

Enrichment was computed across seven annotation databases: Gene Ontology Biological Process, Molecular Function, and Cellular Component (enrichGO with org.Hs.eg.db v3.20.0); KEGG (enrichKEGG, organism hsa); Reactome (ReactomePA v1.50.0 with reactome.db v1.89.0) [37]; and the MSigDB Hallmark and C7 immunologic collections (msigdbr v25.1.1). All analyses used clusterProfiler v4.14.6 [38]. Where CpG-number bias is a concern (enrichment of host genes of off-gene CpGs against an EPIC-array background), enrichment was additionally computed with missMethyl::gometh v1.40.3 [39], which accounts for the unequal number of array probes per gene. Note the specific probe-number-bias-correction method (GOmeth) that gometh implements is more precisely described in a later paper by an overlapping author group (Maksimovic, Oshlack, and Phipson 2021, Genome Biology, PMID 34103055); we cite the original 2016 missMethyl package paper here, consistent with the manuscript's existing reference list, but flag the 2021 paper as an optional additional citation for this specific claim. P-values were Benjamini-Hochberg corrected within each collection independently; enrichment results across collections were not further combined or corrected.

##### 2.20 TRRUST regulatory degree and STRING protein-interaction exploratory analysis

Each gene's transcriptional regulatory context was characterized using TRRUST v2 [32], a manually curated database of human transcription factor to target-gene relationships derived from published literature. Regulatory out-degree (number of downstream curated targets; a proxy for whether the gene is itself a lineage master regulator) and in-degree (number of upstream TF regulators) were related to cross-validated $R^2$ by Pearson correlation. Genes absent from TRRUST v2 were assigned a degree of zero. An initial exploratory analysis used STRING protein-protein interaction network degree as a proxy for regulatory complexity; this produced no association (r approximately 0) and is reported only as a methodological note, because PPI degree does not measure directed transcriptional regulation and is inflated by publication bias toward well-studied proteins.

##### 2.21 PBMC admixture transfer: partial-correlation covariate definitions

T cell models were applied to whole-PBMC methylation from 385 MAG cohort PBMC samples. Methylation-predicted expression from PBMC methylation was compared to each subject's matched purified T cell NanoString expression via Pearson correlation, yielding a PBMC transfer correlation per gene (n = 362 paired samples, all 77 genes). To distinguish a genuine methylation identity signal from a cell-fraction readout (the null hypothesis being that higher predicted scores in PBMC simply track a higher T cell fraction), partial correlations were computed controlling for the PBMC T cell fraction alone and then controlling for the same 7 grouped cell-fraction covariates used in the cancer-cohort analysis above (CD8_total, CD4_total, Treg, B_total, NK, Mono, Neu; cent12CT reference, RPC method; eosinophils and basophils excluded for the same rank-deficiency reason). A signal surviving this full composition adjustment is intrinsic to the gene's methylation pattern rather than a composition proxy.

#### 2.22 PBMC-retrained admixture-aware models: gene panel and parameters

As an independent test, admixture-aware models were retrained directly on PBMC methylation to predict purified T cell expression, for a representative panel of 12 genes (CD8A, KLRG1, GZMB, TBX21, RUNX3, EOMES, CD3E, SPI1, FOXP3, IL6, CD274, and PDCD1), chosen to span robust high-predictability lineage markers, composition-sensitive genes expected to behave differently in an admixed sample (CD3E, SPI1), and additional mid- and low-predictability comparison genes – not all 77 genes. Retraining used the same nested cross-validation procedure as the primary models, but with a single alpha of 0.5 and a single subject-level 80/20 split (seed 42) rather than the full six-value alpha grid and three-fold outer cross-validation used for the primary models; this analysis is therefore exploratory and illustrative of the transfer/retrain comparison, not a definitive optimized PBMC model. PBMC-retrained models were compared to T cell models on matched held-out performance metrics (held-out test $R^2$ and cross-validated $R^2$) for these 12 genes. CpG overlap between T cell and PBMC models was computed as the fraction of T cell model stable CpGs present in the PBMC model's selected feature set (median approximately 1.5% across the 12 genes).

#### 2.23 MGB Biobank whole-blood deployment: scoring, CD80 handling, and covariate scheme

To demonstrate deployment at population scale in the most accessible sample type, both the T cell-weighted and PBMC-weighted model sets were scored on the MGB/Harvard Biobank whole-blood cohort (Illumina EPIC v1; ssNoob normalization; n = 4,250 with complete phenotype and outcome data; 4,483 with age for the validity gate). Scoring used a memory-safe per-model M-value conversion implemented in the deployment bundle (score_beta_matrix.R); each predicted gene score was standardized to mean 0 and unit standard deviation within the analysis set (per-SD scaling), so all reported hazard ratios and odds ratios are per-SD effects. One gene score, CD80, showed near-zero variance in the PBMC-weighted model and was excluded from that model only, leaving 76 evaluable PBMC scores; CD80 was retained in the T cell-weighted model, where its predicted score showed non-degenerate variance, leaving all 77 T cell scores evaluable. Composition was estimated by EpiDISH RPC with the cent12CT reference [43] as above; all disease models below include 11 of the 12 raw cell fractions as covariates (neutrophils dropped as the sum-to-one reference to avoid singularity). Note this whole-blood covariate scheme (11 of 12 raw fractions) differs from the 7-grouped-covariate scheme used in the cancer-cohort and PBMC analyses above; both choices independently avoid rank-deficiency, but by different means (dropping one reference category here, versus grouping small subsets there), and the two schemes should not be assumed identical.

#### 2.24 MGB incident disease outcome definitions

Disease outcome data were obtained from structured EHR records linked to the MGB Biobank. Incident outcomes were defined as a new ICD-coded diagnosis occurring after the methylation collection date, excluding participants with a prevalent diagnosis at baseline. All-cause mortality was ascertained from hospital records and death registration. Twelve incident disease outcomes were tested (Asthma, COPD, Type 2 Diabetes, Coronary Artery Disease, CVD excluding stroke, Stroke, Congestive Heart Failure, Chronic Kidney Disease, Chronic Liver Disease, Cancer, Depression, and Cognitive Deficit), plus all-cause mortality. These outcomes were selected by the MGB collaborators (M.M., J.L.S.) based on availability and event count in the biobank; no pre-specified primary outcome was designated for the MGB analysis, which is treated as a discovery-and-illustration deployment rather than a confirmatory study.

#### 2.25 Cox proportional-hazards model specification (MGB deployment)

For all-cause mortality and each incident disease, composition-adjusted Cox proportional-hazards models were fit using the survival package v3.7.0 [44, 45] with time-to-event as the follow-up from collection date to event or censoring. Cox PH was chosen because it appropriately handles right-censored follow-up, is the standard model for incident disease in biobank EHR cohorts, and yields hazard ratios (HRs) directly interpretable as per-SD effects on the instantaneous event rate. The proportional-hazards assumption was not formally tested at this scale (912 PBMC models and 924 T cell models); violations would attenuate effect estimates toward the null. For each outcome, models were fit three ways: unadjusted (score only), semi-adjusted (score plus age and sex), and fully adjusted (score plus age, sex, and 11 EpiDISH cell fractions); only fully adjusted results are reported in the main text. Cross-sectional (prevalent) associations used logistic regression with the same covariate structure.

#### 2.26 Methylation array normalization pipeline (training and PBMC datasets)

The training (MAG T cell, n = 333) and PBMC (n = 385) datasets were processed together from IDAT files using a reference-anchored functional normalization procedure. Control-probe intensities for each sample were projected into a principal component space (3 PCs) pre-computed from approximately 22,000 whole-blood EPIC v1 samples (the MGB/Harvard and Generation Scotland cohorts), yielding stable normalization coordinates that are not influenced by the small size or composition of the current batch. Functional normalization was then applied using these projected PCs via a modified preprocessFunnormBatch function (based on preprocessFunnorm in minfi) [24] with background correction and dye-bias correction enabled. The normalized betas were subsequently corrected for residual Type I versus Type II probe-type bias using regression on correlated probes (RCP [25]; betaRCP implementation). CpGs with a detection p-value greater than 0.05 (minfi detectionP) were masked to NA. This procedure was applied identically to both the T cell and PBMC datasets; because the PBMC samples were from a single scan-batch generation (206xxx chips only), the global-reference PCA anchoring was the primary source of inter-batch standardization.

##### 2.27 NanoString expression normalization pipeline

T cell gene expression was measured on the NanoString nCounter Human T Cell Characterization Panel (77 genes plus housekeeping controls). Raw counts were upper-quartile normalized to correct for total-count differences across samples, followed by removal of unwanted variation using RUVg (k = 1, using endogenous spike-in probes as negative controls; RUVSeq [27]) to remove systematic technical effects not captured by the housekeeping probes. Size-factor normalization was then applied via DESeq2 [28] and counts were log2-transformed. Two panel targets for which more than 25% of samples fell at the background-expression floor (GRAIL/RNF128 and VTCN1/B7-H4) were excluded as unevaluable, leaving 77 T cell genes for modeling.

##### 2.28 CpG feature-selection strategy comparison

For each of the 77 genes, candidate CpGs were identified genome-wide by methylation-expression association. Three feature-selection strategies were evaluated and selected between by cross-validation: Pearson correlation (computed across all training samples), Spearman correlation (rank-based, robust to outliers), and a limma linear model with duplicateCorrelation to accommodate the repeated-measures structure of longitudinal samples [29, 30]. The duplicateCorrelation approach estimates the within-subject correlation as a nuisance parameter and adjusts the standard errors accordingly, yielding more calibrated association statistics under the paired design. The top 50,000 CpGs by association statistic were retained as candidate features for each gene.

##### 2.29 Stable-CpG definition and cross-validated-$R^2$ performance tiers

Stable CpGs were defined as those with a non-zero elastic-net coefficient in at least two of the three outer folds, requiring consistency of selection across data partitions rather than relying on a single fit. Genes were grouped into four interpretive performance tiers by cross-validated $R^2$ (High at or above 0.70; Moderate 0.55 to 0.70; Weak 0.40 to 0.55; Uncoupled below 0.40); these boundaries are an interpretive convention applied to a continuous distribution, and all quantitative analyses use $R^2$ as a continuous variable.

##### 2.30 Multiple-testing correction: MGB deployment test families

Benjamini-Hochberg false discovery rate correction was applied separately within each model set, with a significance threshold of FDR less than 0.05: 912 gene-disease tests (76 genes times

12 diseases) for the PBMC-weighted model, and 924 gene-disease tests (77 genes times 12 diseases) for the T cell-weighted model. Separate FDR correction for mortality was applied independently (76 tests for the PBMC-weighted model, 77 tests for the T cell-weighted model). No correction was applied across the PBMC and T cell model sets; the two models were treated as parallel discovery analyses rather than replication.

##### 2.31 T cell/PBMC model concordance methodology

To characterize the concordance and complementarity of the two model sets, Pearson correlation of log(OR) was computed across all 912 gene-disease incidence pairs for the 76 genes common to both model sets (CD80, evaluable only in the T cell-weighted model, was not included in this comparison). Low overall concordance ($r = 0.01$) indicates complementary rather than redundant readouts, consistent with the 1.5% CpG overlap between T cell and PBMC-retrained models established in the admixture analysis (12-gene panel; see PBMC admixture transfer, above).
